# Inhibiting TG2 sensitizes lung cancer to radiotherapy through interfering TOPOIIα-mediated DNA repair

**DOI:** 10.1101/597112

**Authors:** Xiao Lei, Zhe Liu, Kun Cao, Yuanyuan Chen, Jianming Cai, Fu Gao, Yanyong Yang

**Author notes:** Authors contributed equally to this work (Xiao Lei, Zhe Liu, Kun Cao, Yuanyuan Chen). Corresponding author: Yanyong Yang, Fu Gao and Jianming Cai. Address: Department of Radiation Medicine, Faculty of Naval Medicine, Second Military Medical University; 800, Xiangyin Road, 200433, Shanghai, P.R. China.

## Abstract

Radiotherapy is an indispensable strategy for lung cancer, however, treatment failure or reoccurrence is often found in patients due to the developing radioresistance. Novel approaches are required for radiosensitizing to improve the therapeutic efficacy. In present study, we found that transglutaminase 2 (TG2) confers radioresistance in non-small cell lung cancer (NSCLC) cells through regulating TOPOIIα and promoting DNA repair. Our data showed that TG2 inhibitor or knockdown increased NSCLC radiosensitivity in vivo and in vitro. We found that TG2 translocated into nucleus and located to DSB sites, surprisingly, knockdown TG2 or glucosamine inhibited the phosphorylation of ATM, ATR and DNA-Pkcs. Through IP-MS assay and functional experiments, we identified that TOPOIIα as an downstream factor of TG2. Moreover, we found that TGase domain account for the interaction with TOPOIIα. Finally, we found that TG2 expression was correlated with poor survival in lung adenocarcinoma instead of squamous cell carcinoma. In conclusion, we demonstrated that inhibiting TG2 sensitize NSCLC to IR through interfere TOPOIIα mediated DNA repair, suggesting TG2 as a potential radiosensitizing target in NSCLC.

## Introduction

Radiotherapy is an indispensable strategy in treating lung cancer, of which 80% is non-small cell lung cancer (NSCLC) with poor outcomes (1, 2). Despite the advance in physical techniques, novel approaches in radiosensitizing from biological aspect are required to overcome the growing radioresistance during radiotherapy. The current research involving radiosensitization mainly falls in the following fields: DNA damage repair, poly (adenosine diphosphate–ribose) polymerase inhibitors, histone deacetylase inhibitors, tumor hypoxia and redox conditions, antiangiogenic drugs *etc* (3). However, most of these drugs are in research process or clinical trials, efficacy as well as normal tissue toxicity limits their application.

Transglutaminase 2 (TG2), a member of Transglutaminases family, exerts multiple physiological functions and is associated with cancer cell survival, metastatic behavior and chemoresistance (4–7). It has been proved that TG2 was related to multiple drug resistance including cisplatin, histone deacetylase inhibitor, EGFR-TKI *etc* (5, 8, 9). Recently, the prognostic value of elevated TG2 for patient survival has been illustrated in NSCLC and are attracting more and more attention (10, 11). These studies indicated that TG2 might be critical for radiation resistance in NSCLC. When we are preparing this manuscript, Sheng *et al*. reported that TG2 inhibitor KCC009 induces radiosensitization in lung adenocarcinoma cells(12). However, the detailed role of TG2 in NSCLC radioresistance and the underlying mechanism remains unclear.

Previous studies indicated that TG2 was related to DNA damage repair, which is aberrant active in cancer cells (13–15). Previous study had showed that ATM inhibitor KU55933 abrogated the constitutively activation of TG2 induced by genotoxic drug MNNG (13). ATM mediated NF-kB activation increased the level of TG2. TG2 was also proved as a target of p53 and involved in DNA damage repair, and knockdown of p53 reduced the level of TG2 (14, 15). But the response of TG2 to ionizing radiation and the exact role of TG2 in DNA repair remains to be uncovered.

Here, we report that TG2 confers to radioresistance in NSCLC and enhanced DNA repair capacity through directly interacting with DNA topoisomerase IIα (TOPOIIα). We found that ionizing radiation (IR) resulted in a rapid nuclear translocation of TG2 and knockdown TG2 significantly inhibited DNA repair. TG2 was found to bind and activate TOPOIIα in nucleus to initial DNA damage repair processes, such as phosphorylation of ATM, ATR and DNA-PKcs. Moreover, we used a clinically used TG2 inhibitor, glucosamine, and found it significantly sensitized lung cancer to IR in vivo and in vitro. Finally, we found TG2 was significantly correlated with the survival in lung adenocarcinoma instead of squamous cell carcinoma patients, which suggest possible prognostic value of TG2 in lung adenocarcinoma. These data provide the possibility of clinical translation of TG2 inhibitor in the radiosensitization of lung cancer.

## Results

### Inhibition of TG2 sensitizes lung cancer cells to ionizing radiation

It has been proved that TG2 high expression was related to chemoresistance of multiple cancers (16–18). To determine whether TG2 participates in radioresistance in NSCLC, firstly we used a TG2 inhibitor, glucosamine, which was already used in clinics as an anti-inflammatory drug. We found that glucosamine effectively reduced cell viability in A549 cells, while showed little influence on normal lung BEAS-2B cells (Fig. 1A). Besides, glucosamine effectively inhibited TG2 level at the concentration of 5mM in A549 cells (Fig. 1B). Compared with normal lung BEAS-2B cells, TG2 expression was also found to be elevated in lung adenocarcinoma cell lines including A549, H1975 and H358 cells. (Fig. 1C, S1A). By using colony formation assay, we found that glucosamine or TG2 knockdown significantly sensitized A549, H1299, H460 cells to IR, while glucosamine showed no further sensitizing effects on TG2 knockdown cells (Fig. 1D-F). This data was also confirmed in CRISPR Cas9 mediated TG2 knockout cells (Fig. 1G). Alternatively, we used apoptosis assay to determine cellular damage in TG2 inhibited cells. It was found that glucosamine treatment resulted in more apoptotic cells in response to IR, while glucosamine showed no sensitizing effects on BEAS-2B cells (Fig. 1H, Fig. S1B, C).

**Figure1.**
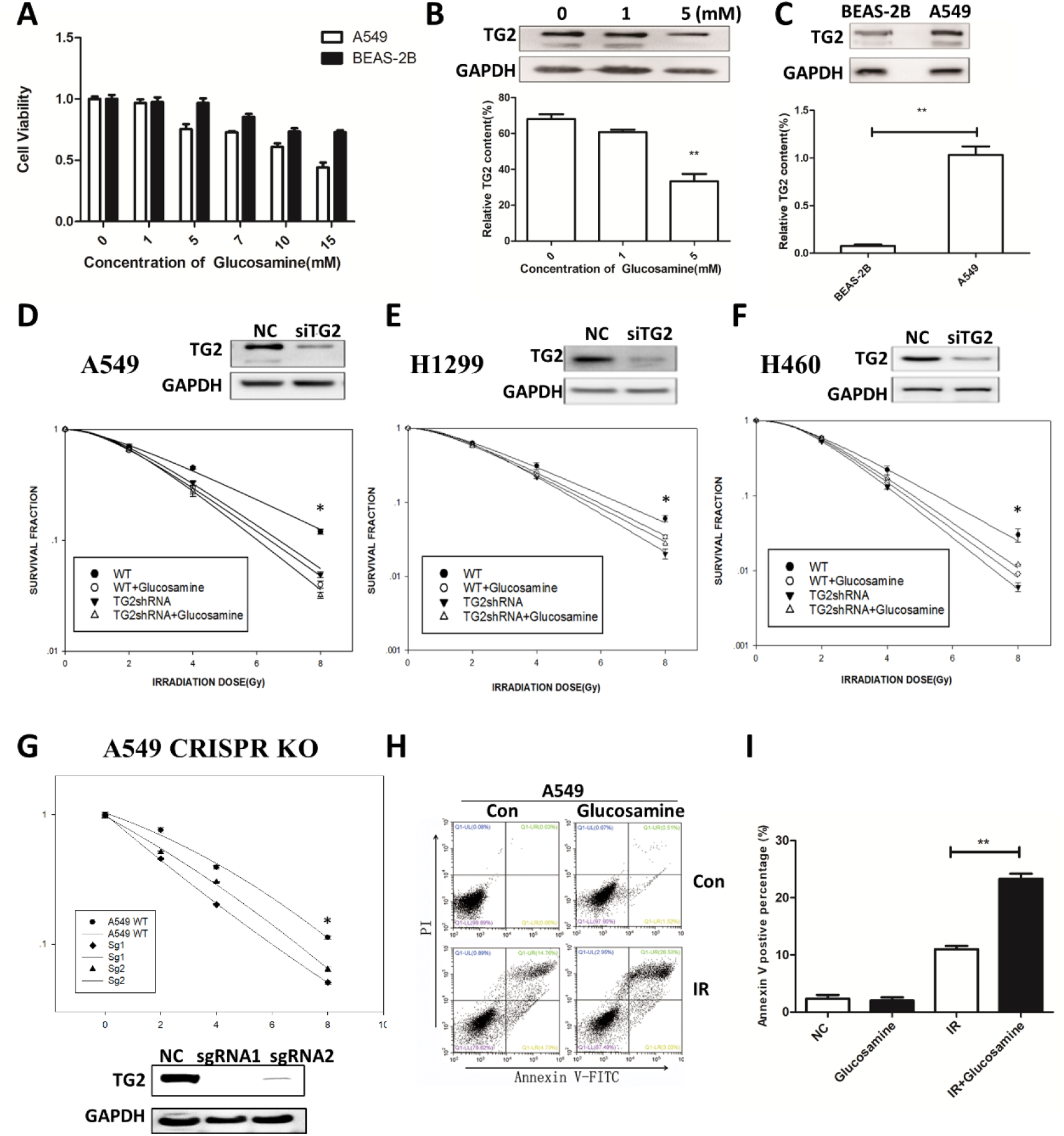
Inhibition of TG2 sensitize lung cancer cells to ionizing radiation. (A)A549 and BEAS-2B cell lines were analyzed for their cell viability after treated with different concentration of glucosamine. (B) Expression of TG2 in A549 cells with 0, 1, 5mM glucosamine pretreated. (C) Expression of TG2 in A549 and BEAS-2B cells. (D-F) A549, H1299, H460 and their TG2 knock down cell lines were analyzed for their colony forming ability against IR with/without glucosamine (5mM) pretreated. (G) A549 WT and CRISPR KO cells were analyzed for their colony forming ability against IR. (H, I) Flow cytometric analysis of A549 cell line against 8Gy irradiation with/without glucosamine (5mM) pretreated. *P < 0.05, **P<0.01 versus radiation group.

### Radiation induces TG2 nuclear translocation and initiates DNA damage response

To figure out how TG2 confers to radioresistance, we investigated its subcellular location and the relationship with DNA damage repair, the main effects of radiation response. By using Immunofluorescence staining and nuclear protein western blot assay, we found that radiation rapidly induced TG2 nuclear translocation, which could be inhibited by glucosamine (Fig. 2A, B, S2B). Based on distinct functions of TG2, we used calcium inhibitor perillyl alcohol (POH), TG2 activity inhibitor cystamine, and NF-kB inhibitor QNZ, our data showed that QNZ inhibited radiation-induced nuclear translocation of TG2 (Fig. S2A). Moreover, we found that glucosamine treatment significantly inhibited the phosphorylation of DNA-PKcs, ATM and ATR, which are critical for initiating DNA damage repair (Fig. 2C). Further, we used a siRNA of TG2 to investigate its role in DDR, and found the same effects on DNA-PKcs, ATM and ATR inhibition. However, TG2 siRNA combined with glucosamine treatment didn’t showed any additive effects (Fig. 2D). To investigate the influence of TG2 inhibitor on DNA damage, we examined γH2AX foci and found that TG2 knockout significantly impaired DNA repair in response to IR (Fig. 2E, F). By using a comet assay, we confirmed that more DNA damage remains unrepaired in cells treated with glucosamine (Fig. 2G, H, I).

**Figure2.**
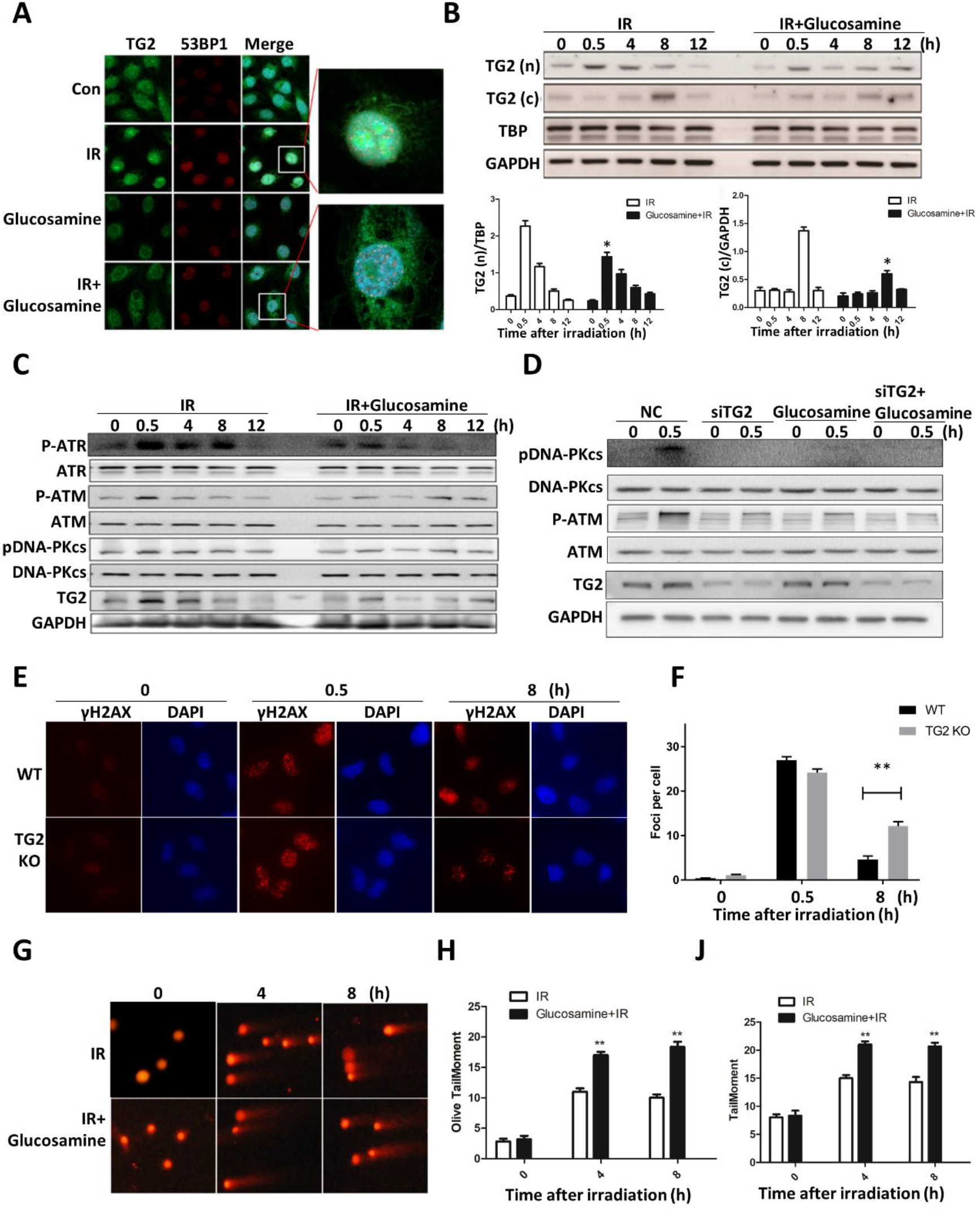
Radiation induces TG2 nuclear translocation and initiates DNA damage response. (A) Immunofluorescence of TG2 and 53BP1 in A549 cells exposed to IR with/without glucosamine (5mM) pretreated. (B) Immunoblot of endogenous TG2 in the cytoplasmic and nuclear fractions of A549 cells exposed to 8Gy irradiation with/without glucosamine (5mM) pretreated. (C)A549 cells were exposed to IR with/without glucosamine (5mM) pretreated and harvested at the indicated time points. Whole cell lysates were analyzed with indicated antibodies. (D)A549 and A549 TG2 knock down cells were exposed to IR with/without glucosamine (5mM) pretreatments and harvested at 0, 0.5h. Then whole cell lysates were analyzed with indicated antibodies. (E) A549 and A549 TG2 KO cells exposed to IR were immunofluorescent stained against γ-H2AX (green) and DAPI (blue). (F) The average numbers of γ-H2AX foci per cell among A549 and A549 TG2 KO cells exposed to 2Gy irradiation. (G) Representative comet assay showing the tail moment of A549 cells exposed to IR with/without glucosamine (5mM). n = 3 independent experiments. Quantification in H and J; data represent mean ± SEM. *P < 0.05, **P < 0.01 versus radiation group.

### TG2 interacts with TOPOIIα and participates in DNA repair

To identify the specific target of TG2, we conducted an Immunoprecipitation–Mass Spectrometry (IP-MS) assay in A549 cells. Through bioinformatics analysis, we found that 134 proteins bind to TG2 in radiation group compared with normal group (Fig. 3A, Table S1). Among these, TOPOIIα was found to be related to DNA damage, and TOPOII inhibitors were clinically used in cancer therapy, such as etoposide and doxorubicin. Then we used lazer assay and found that after lazer irradiation both TG2 and TOPOIIα were recruited in DSB site (Fig. 3B, Fig. S3A). Then we used immunoprecipitation assay and proved that TG2 bind to TOPOIIα after IR (Fig. 3C), as a consequence of its nuclear translocation. To confirm the direct binding of these two proteins, we transfected TG2 and TOPOIIα into 293T cells. The interaction of TG2 and TOPOIIα was further confirmed in co-immunoprecipitation experiments (Fig. 3D, E). Functionally, knockdown of TOPOIIα resulted in more DNA damage after IR (Fig. 3F). Moreover, TOPOIIα knockdown together with glucosamine didn’t show additive effects on cellular DNA damage (Fig. 3F). TG2 knockdown also causes more γH2AX accumulation in TG2 knockdown or TOPOIIα knockdown cells after irradiation (Fig. S3C). Then we investigated the influence of TOPOIIα on cellular radiosensitivity, and found that TOPOIIα knockdown cells are more sensitive to radiation-induced cell death (Fig. S3B). However, no significant difference was found in TOPOIIα knockdown cells compared with TOPOIIα-TG2 double knockdown. To determine whether TOPOIIα participate in DNA damage repair, we used a siRNA and found that TOPOIIα siRNA inhibited radiation-induced phosphorylation of DNA-PKcs as well as ATM (Fig. 3G). And similar results were observed in TOPOIIα knockdown cells together with TG2 knockdown or glucosamine treatment (Fig. 3G). Then we transfected TOPOIIα into TG2 knockdown cells, and found that TOPOIIα overexpression rescued the inhibitory effects of TG2 knockdown on DDR pathway (Fig. 3G). However, TG2 overexpressing in TOPOIIα knockdown cells showed no influence on DNA damage response (Fig. 3H).

**Figure3.**
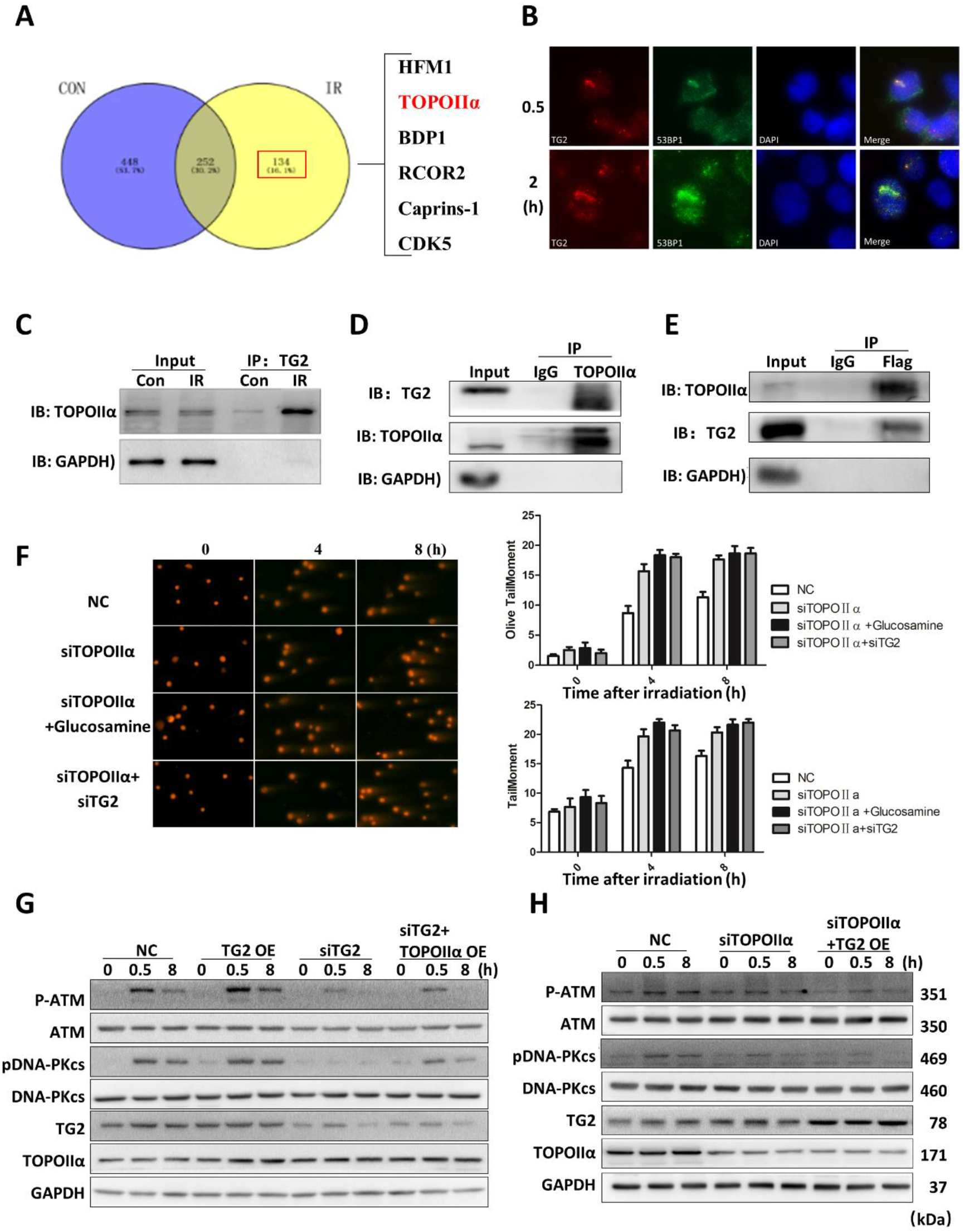
TG2 interacts with TOPOIIα and participate in DNA repair. (A) Venn diagram about bioinformatics analysis in Immunoprecipitation–Mass Spectrometry (IP-MS) assay in A549 cells. (B) Representative immunofluorescence of TG2 and 53BP1 at DNA damage sites, in HT1080 cells following laser microirradiation. (n = 3 independent experiments). (C)A549 cells exposed to IR were harvested at 0, 0.5h. Whole cell lysates were subjected to IP with anti-TG2 antibody and immunoblotted with TOPOIIα antibody. (D) 293T cells transfected with FLAG-tagged TG2 and TOPOIIα constructs were exposed to IR (8Gy) and harvested for 30mins. Whole cell lysates were subjected to co-IP with anti-TOPOIIα antibody and western blotted against anti-TG2 and anti-TOPOIIα antibodies. (E) 293T cells transfected with FLAG-tagged TG2 and TOPOIIα constructs were exposed to IR (8Gy) and harvested for 30mins. Whole cell lysates were subjected to co-IP with anti-FLAG antibody and western blotted against anti-TOPOIIα and anti-TG2 antibodies. (F) Representative comet assay showing the tail moment of A549 cells transfected with TOPOIIα siRNA were exposed to IR with/without glucosamine (5mM) or TG2siRNA. (n = 3 independent experiments). (G) A549 cells transfected with TG2 construct/ TG2 siRNA/ TG2 siRNA and TOPOIIα construct were treated with IR (8Gy) and harvested at the indicated time points. Whole cell lysates were analyzed with indicated antibodies. (H) A549 cells transfected with TOPOIIα siRNA/ TOPOIIα siRNA and TG2 construct were treated with IR (8Gy) and harvested at the indicated time points. Whole cell lysates were analyzed with indicated antibodies.

### TGase domain confers to the interaction with TOPOIIα and radioresistance of NSCLC

TG2 is a multifunctional enzyme with different domains, structurally including four domains: an NH2-terminal β-sandwich domain; a catalytic core domain containing a catalytic triad for the acyl-transfer reaction (Cys277, His335 and Asp358) for acyl-transfer reaction; a β-barrel1 domain, containing GDP/GTP-interacting residues, that is involved in receptor signaling and a β-barrel2 domain (19–21). To determine which domain was indispensable for the interaction with TOPOIIα and confers to radioresistance, we generated constructs with different fragments, as well as constructs with mutation for dysfunction of distinct domain (Fig. 4A, B; Fig. S4A). We transfected these fragments or mutations into TG2 low expressing H1299 cells, and determined the cellular radiosensitivity based on survival fraction. Our data showed that TG2 full length expression increased radioresistance, which was only found in ABC domain expression of all fragments (Fig. 4C-D; Fig. S4B). However, in TG2 mutants transfected cells, only W241A mutant showed no increase on radioresistance (Fig. 4E, F; Fig. S4C). Then we performed immunoprecipitation assay in 293T cells transfected with both TG2 fragment or mutant and TOPOIIα. It was found that none of TG2 fragment bind with TOPOIIα efficiently (Fig. 4G), showing that the interaction of TG2 and TOPOIIα might require the cooperation of multiple domains. Then we used TG2 expression vector with different functional mutations, and found that W241A mutant reduced the binding efficacy of TG2 and TOPOIIα (Fig. 4H). Taken together, our data showed that TGase function might be critical for the role of TG2 in radioresistance.

**Figure 4.**
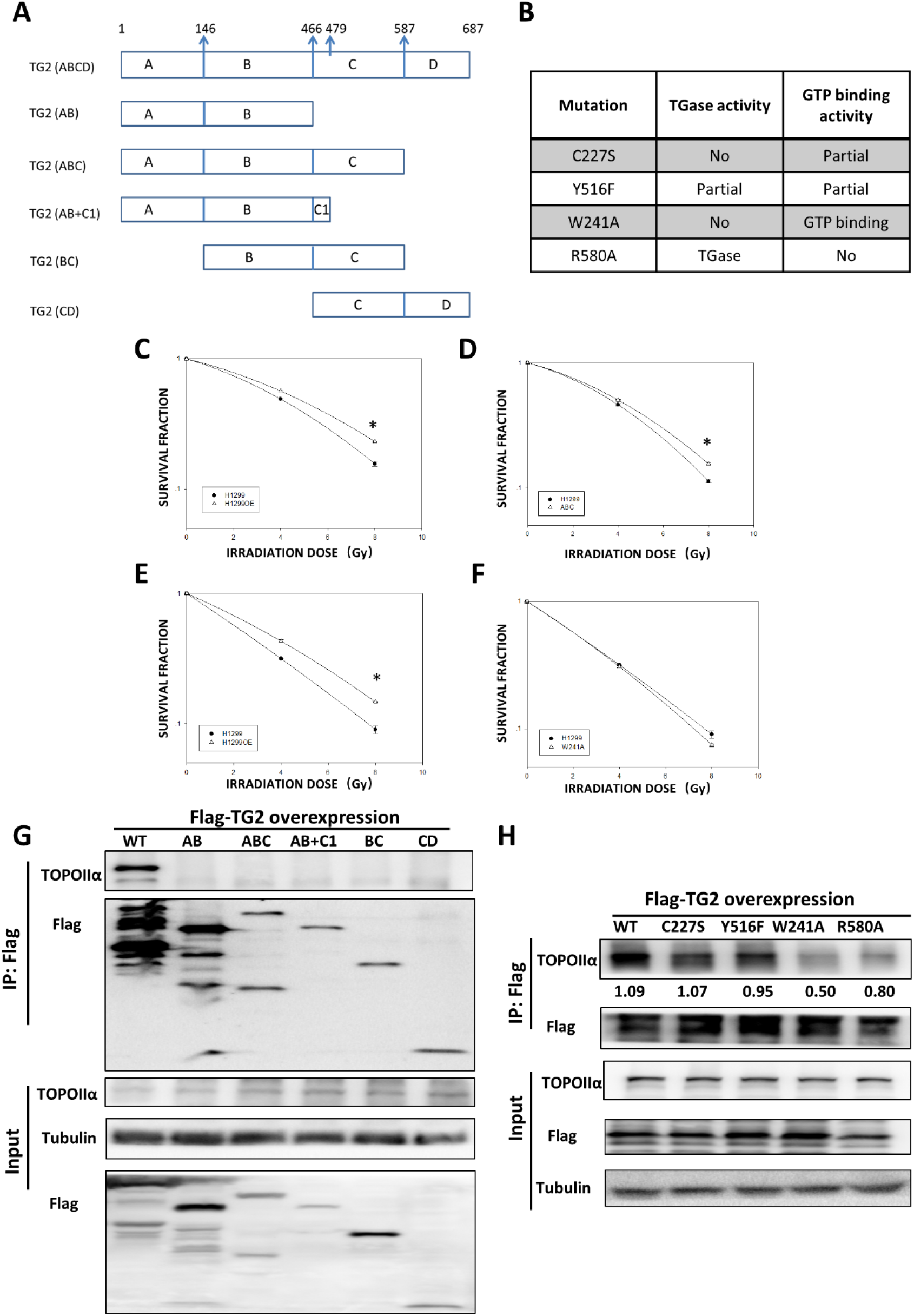
TGase function confers the interaction with TOPOIIα and radioresistance of NSCLC. (A) Schematic structure of TG2 full length, and fragments including AB, ABC, AB+C, B+C, CD. (B) Plasmids encoding wild-type TGM2, W241A, C277S, R580A, Y516F cloned in pLenO-GTP were constructed by Biolink biotechnology(Shanghai) Co.,Ltd. (C-D) Survival assay of wild type TG2 and ABC fragment in response to different doses of radiation. (E-F) Analyzing TG2 mutants transfected cells for their colony forming ability against IR (WT and W241A). (I) immunoprecipitation assay in 293T cells transfected with both TG2 fragment and TOPOIIα. (J) immunoprecipitation assay in 293T cells transfected with both TG2 mutant and TOPOIIα.

### TG2 inhibition sensitizes lung tumor to IR in vivo

To investigate the radiosensitizing effects of TG2 inhibition on NSCLC in vivo, we used an in situ lung cancer model generated by our group, together with local irradiation (Fig. S5A, B and C). It was found that if left untreated, all mice died within one month, while glucosamine treatment significantly increased survival of tumor bearing mice compared with single radiation group (Fig. 5A, B). The tumor size was reduced in glucosamine and radiation treated group (Fig. 5C). Through scanning the largest cross section of tumor, we found that the overall area was also significantly reduced in glucosamine combined with radiation group (Fig. 5D). HE staining of lung tissues observed that glucosamine treatment combined with radiation significantly reduced the size of lung cancer. And no invasion was observed in glucosamine treated group (Fig. 5D, E). Epithelial–mesenchymal transition (EMT) is an important process in the initiation of cancer metastasis. We performed IHC staining of tumor sample and found that glucosamine treatment reduced the level of Vimentin and αSMA, while elevated the expression of E-cadherin (Fig. 5F, G). These data indicated that glucosamine inhibited EMT process, which might accounts for the inhibition of cancer metastasis. TG2 is also found to participate in NF-kB activation. We also found that glucosamine reduced Ki67 and p65 positive cells in tumor area (Fig. 5F, G).

**Figure 5.**
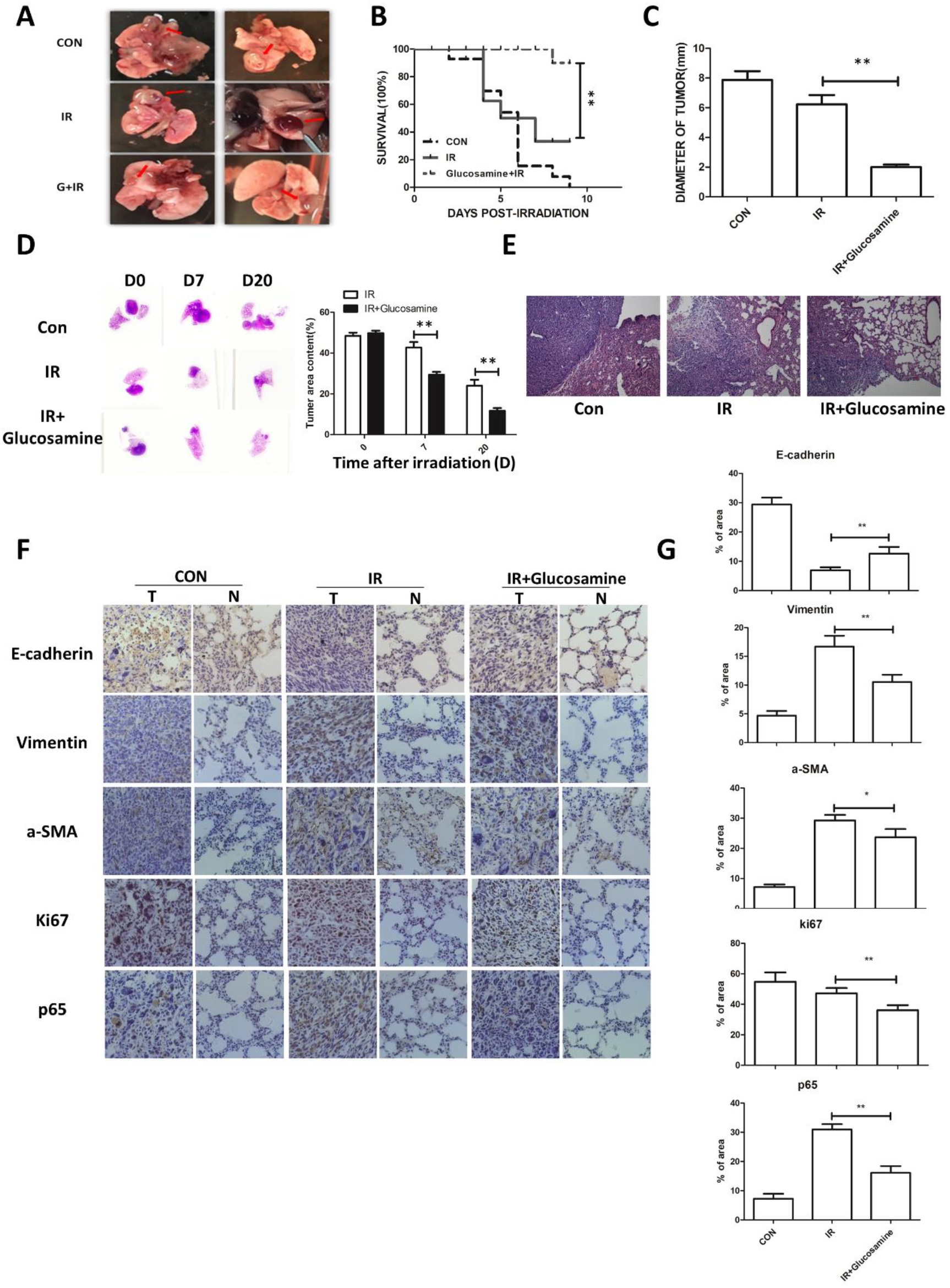
TG2 inhibition sensitize lung cancer to IR in vivo. (A) Tumors formed in female C57BL/6 mice by injection of LLC cells. The recipient mice were treated with whole lung irradiation (15Gy) with/without glucosamine (150 mg/kg/d) for 3 days before IR. (B) Survival rate of tumor bearing mice treated with whole lung irradiation (15Gy) with/without glucosamine (150 mg/kg/d) for 3 days before IR. (C) The mean diameter of each tumor in different groups. (D) the images of each group scanning the largest cross section of tumor in different days. (E) HE staining of lung tissues with tumor in each group. (F)E-cadherin, SMA, Vimentin, Ki67 and p65 staining of lung tissues with tumor after IR with/without glucosamine treated.(G) Quantification of protein expression levels of E-cadherin, SMA, Vimentin, Ki67 and p65 staining of lung tissues with tumor after IR with/without glucosamine treated. Values are given as mean± SEM (n=10), *P<0.05 and **P<0.01 versus single radiation group.

### High expression of TG2 predicts poor survival in lung adenocarcinoma instead of squamous cell carcinoma

It has been reported that TG2 was elevated in NSCLC patient tissues, and high expression of TG2 predicts poor outcome of disease free survival as well as overall survival (11). TG2 expression was also found to be related to survival of patients treated with EGFR-TKIs (22). However, we collected 80 pairs of tissues from NSCLC patients and also found that in some patients, TG2 mRNA expression was downregulated in lung cancer tissues compared with adjacent normal lung tissues (Fig. 6A). There is no evidence showing that TG2 expression was related to clinical features of age, gender and family cancer history (Table S2). When referred to cancer pathological information, we found that TG2 was mainly highly expressed in glandular tubular adenocarcinoma (Figure 6A). However, TOPOIIα expression was not consistent with TG2, which indicates that maybe other mechanism is involved (Fig. 6B). TG2 protein expression was examined with IHC assay (Figure 6C, D). Then the association between TG2 expression and patient survival was examined with a Kaplan-Meier plotter tool (www.kmplot.com). Our data showed that high expression of TG2 was not associated with overall survival in all set of patients (Fig. S7, P=0.0074, dataset 216183; P=0.075, data set 211573; P=0.19, dataset 211003). Surprisingly, high TG2 expression was significantly associated with overall survival in lung adenocarcinoma lung cancer patients in three dataset (Fig. 6F to H, P=0.0035, dataset 216183; P=4.7e-06, data set 211573; P=0.00056, dataset 211003). While no significant difference was found in squamous cell carcinoma (Fig. S7, P=0.61, dataset 216183; P=0.37, data set 211573; P=0.34, dataset 211003). When compared with radiotherapy, no significant difference was found in patient with high TG2 or low TG2 expression (Fig. S8). With small sample size, we could still see the trends showing the possible association of TG2 and patients outcomes.

**Figure6.**
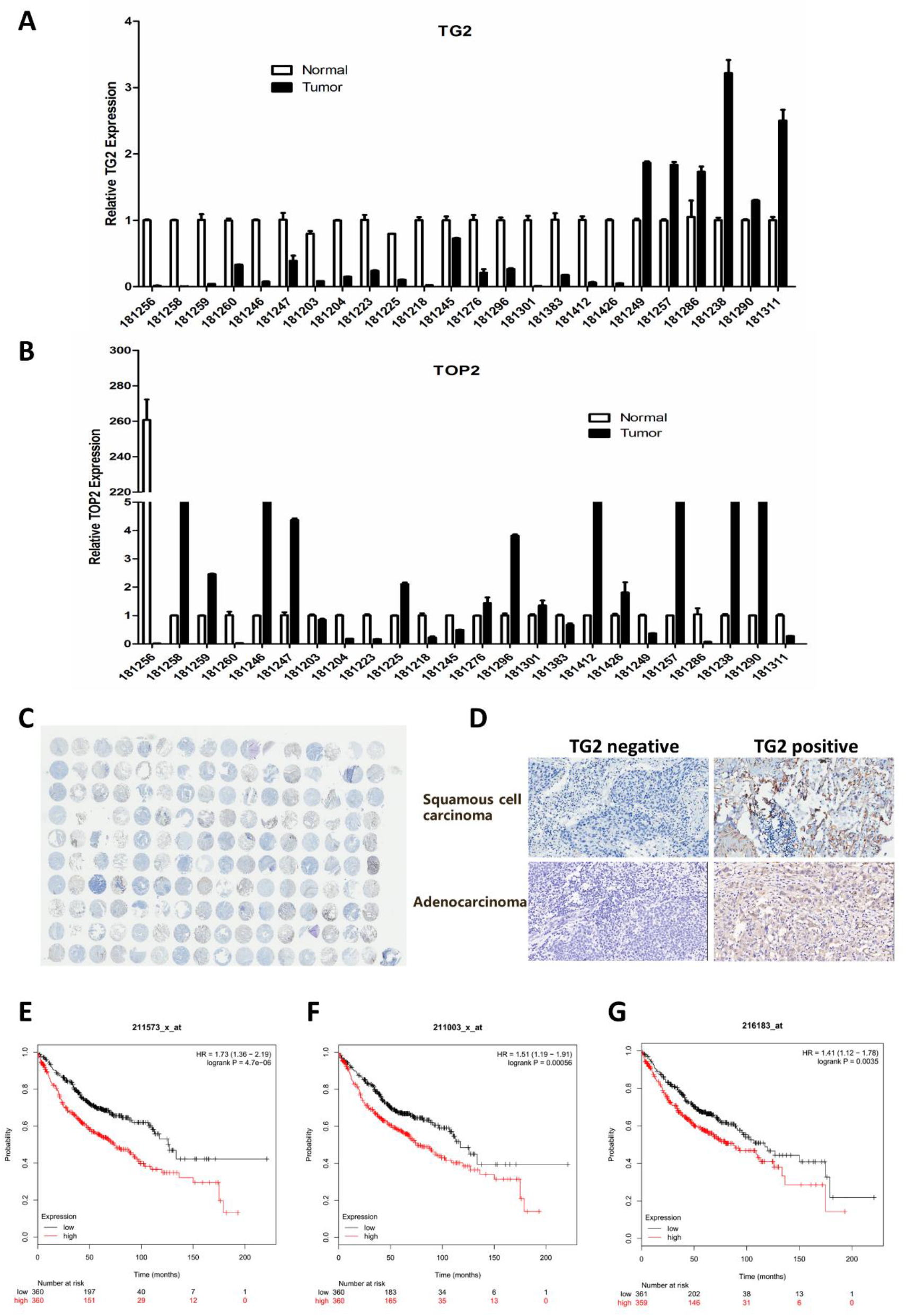
High expression of TG2 predicts poor survival in lung adenocarcinoma instead of squamous cell carcinoma. (A) TG2 mRNA expression in lung cancer tissues compared with adjacent normal lung tissues. (B) TOPOIIα level in lung cancer tissues compared with adjacent normal lung tissues. (C) scanning of tissue array of the samples we collected from NSCLC patients. (D) representative images of TG2 negative and positive staining of lung cancer samples. (E-G) Kaplan-Meier survival curve showing significant association between TG2 IHC staining and overall survival of patients from lung adenocarcinoma from three set of microarray (P=0.0035, dataset 216183; P=4.7e-06, data set 211573; P=0.00056, dataset 211003).

**Figure 7.**
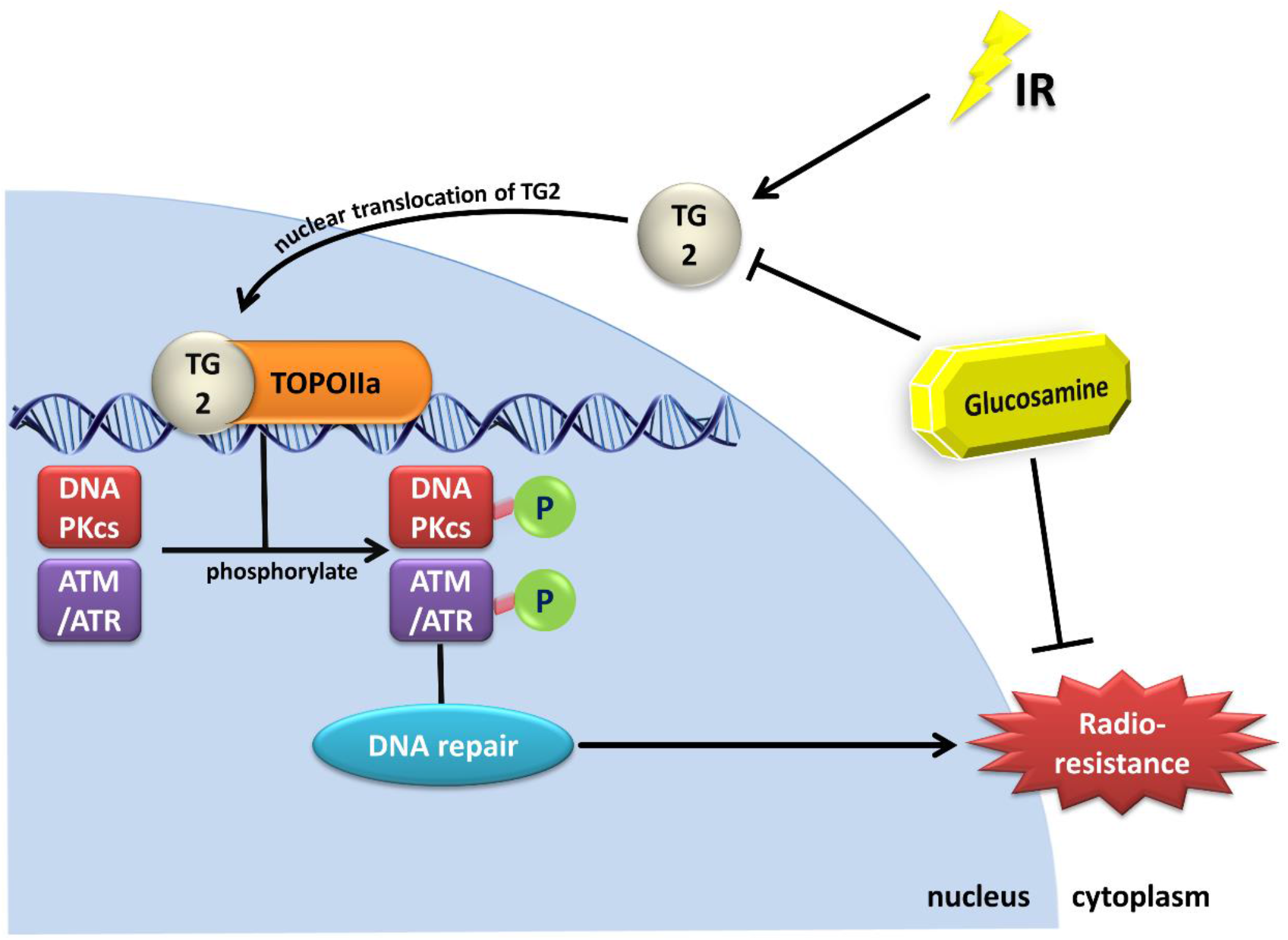
Proposed model illustrating the mechanism how TG2 confers to radioresistance. In response to radiation, TG2 translocate into nucleus and is recruited to DSB sites. The recruitment of TG2 initiate phosphorylation of DNA-PKcs and ATM through interacting with TOPOIIα, which promoted DNA repair. After glucosamine treatment or TG2 knockdown, TG2-TOPOIIα mediated DNA repair was abrogated, and the cancer cells were radiosensitized.

## Discussion

In this study, we demonstrated that TG2 confers to radioresistance in NSCLC through promoting DNA repair. We found that TG2 inhibitor glucosamine significantly sensitize cancer cells and in vivo tumor to radiotherapy. For the first time, we observed significant nuclear translocation of TG2 in response to IR, and we found that TG2 knockdown abolished the activation of DDR signaling pathway. To identify the specific target of TG2, we used an IP-MS method and found that TG2 directly interact with TOP2 and promoted DNA repair, which might account for radioresistance. Finally, we identified that TGase catalytic function of TG2 was critical for DNA repair and radioresistance in NSCLC. Our data provide novel insight for the targeting TG2 in radiosensitization of lung cancer.

### Targeting TG2 for radiosensitization in NSCLC

TG2 is ubiquitously expressed in many tissues and participate in multiple physiological functions, including wound healing, cancer metastasis, apoptosis as well as cell adhesion (23, 24). Recently, the increase level of TG2 was also shown was also related to chemoresistence in cancer, and targeting TG2 provide possibility of overcoming drug resistance (11, 25). In the present study, we found that TG2 inhibitor and siRNA significantly increased radiosensitivity of lung cancer cells. Then through an in vivo lung cancer model, we also proved that glucosamine sensitize lung cancer to IR. These findings indicate that TG2 contribute to resistance to drug or radiation induced cell death. However, it has been proved that TG2 play diverse roles in regulating cell death. On one hand, TG2 promote cell survival through activation of NF-kB without degradation of I-kB, and also induces cell resistance to chemotherapy (26). TG2 also promote chemoresistance through Akt activation, and it also inhibited cell apoptosis through downregulating Bax (27). On another hand, it was shown that TG2 promotes caspase dependent and independent apoptosis through Calpain/Bax Protein Signaling Pathway (28). Besides apoptosis and NF-kB signaling pathway, it was found that the main mechanism of drug resistance was related to cathepsin D, nucleophosmin depletion through the crosslink activity (29, 30). However, targeting TG2 for radiosensitization and its underlying mechanism was largely unknown.

### Radiation induces TG2 translocation to nuclear and participate in DDR

On normal status, TG2 is predominantly located in cytoplasm, even there is some located in the nucleus, the mitochondria, on the plasma membrane, or in the extracellular cell surface (31, 32). Extracellular TG2 is mainly involved in wound healing and scarring, tissue fibrosis and cancer metastasis (33). And under some stimuli, TG2 was shown to translocate to nucleus. In hepatocellular carcinoma cells, acyclic retinoid induced significant nuclear translocation of TG2 and promoted cell death (34). To our knowledge, it is the first time that we observed nuclear translocation of TG2 in response to ionizing radiation, although the significance of TG2 nuclear localization had also been illustrated in other studies. To determine the reason of TG2 nuclear translocation, we used calcium blocker, NF-kB inhibitor, TG2 inhibitor etc, and found that NF-kB might play a critical role in this process.

Then by using TG2 inhibitor and siRNA, we found that TG2 inhibition impaired the activation of DNA repair pathway, including both NHEJ and HR, which is key mechanism for radiation response (35, 36). We found that the phosphorylation of ATM, DNA-PKcs, ATR were all suppressed, while more unrepaired DNA damage was observed. These data indicate that TG2 confers to DNA repair, the elevation of which promote radioresistance. It has been proved that TG2 is related to p53 and ATM function, but our data showed an earlier role of TG2 in DNA damage response, as TG2 tanslocate into nucleus at several minutes after irradiation. Through a lazer assay, we observed that TG2 as well as TOPOIIα were recruited to the DSB site, which suggest TG2 as a direct regulator of DNA repair. However, the exact role of TG2 in DNA repair was to be uncovered.

### Interaction of TG2 and TOPOIIα in DDR response

The significance of TG2 in many pathological processes has drawn much attention, and it has been proved that TG2 regulating many key proteins, including p65, p53 (37–39). To study its role in DNA repair, we performed an IP-MS analysis and found several potential interacting proteins. Among these, we identified TOPOIIα as a target of TG2 and proved their interaction through IP assay. Surprisingly, we found that TOPOIIα knockdown inhibited activation of DNA repair pathway, and TG2 and TOPOIIα double knockdown didn’t produce an additive effect. And for many years, multiple TOPOIIα inhibitors, such as etoposide, doxorubicin, have been used clinically to treating cancer, as well as in radiosensitization (40–42). Thus, inhibition of TG2-TOPOIIα signaling might account for the radiosensitizing effects of TG2 inhibitors. Then based on the distinct function and domain of TG2, we constructed several plasmid expressing different domain or different mutation of TG2. and we found that the TGase function of TG2 was indispensable for its interaction with TOPOIIα and accounts for radioresistance. Our data provide novel mechanism of TG2 in radioresistance. The TGase domain was also required for radioresistance.

Taken together, our findings suggest TG2 as a potential radiosensitizing target for lung cancer radiosensitization in vivo and in vitro. We also provide novel mechanism for the role of TG2 in radioresistance: after irradiation, TG2 translocated into nucleus, bind to DSB site, and initiate DNA damage repair through interacting with TOPOIIα. TG2 was correlated with outcomes of lung adenocarcinoma, and glucosamine provide possibility of translation in clinical applications.

## Materials and methods

### Reagents and plasmids

Glucosamine was purchased from Sigma.The following antibodies were used: anti-TG2 (Abcam, US; 1:1000), anti-Flag (Abcam, US; 1:1000), anti-γ-H2AX (Abcam, US; 1:1000), anti-Rad51(Abcam, US; 1:1000), anti-P-ATM, anti-ATM(Abcam, US; 1:1000), anti-pT2069-DNA-PKcs, anti-DNA-PKcs (Abcam, US; 1:1000), anti-P-ATR, antiATR (Abcam, US; 1:1000), anti-TOPOIIα (Abcam, US; 1:1000), anti-TBP, anti-GAPDH (sampler kit from Cell signaling technology, US; 1:1000), horseradish peroxidase (HRP)-conjugated anti-rabbit or anti-mouse IgG (all purchased from Cell Signaling Technology). Plasmids encoding different TGM2 fragments plasmids and different mutants (C277S, W241A, R580A, Y516F), cloned in pLenO-GTP were constructed by Biolink biotechnology (Shanghai) Co.,Ltd.

### Animals and glucosamine treatments

The whole protocols were approved by the Ethics Committee of Second Military Medical University, China. Female C57BL/6 mice, 8 weeks old, obtained from the Experimental Animal Center of Chinese Academy of Sciences, Shanghai, China, were used for the animal experiment. Mice were fed in daily-changed individual cages, at 25±1°C with food and water provided for free access. All of the animals were implanted with mouse Lewis lung cancer (LLC) cells (25ul, 2×10^5^cells) in the fixed location which was 5mm distance from the lower end of the xiphoid process and parallel to it in right lung, the needle inserted depths was 5mm. The treated mice were randomly divided into four groups: group 1, non-irradiated +saline control; group 2, irradiation + saline; group 3, irradiation + glucosamine. Either glucosamine (150 mg/kg/d) was delivered to the corresponding groups by intraperitoneal injection 3 days before expose to whole lung irradiation.

### Cell culture and glucosamine treatments

Mouse lewis lung cancer (LLC), human NSCLC (A549, H460, H1299, H1975, H358) and human bronchial epithelial cell line BEAS-2B were purchased from the ATCC (USA). YFP-53BP1-HT1080 was provided by Division of Molecular Radiation Biology, Department of Radiation Oncology, University of Texas Southwestern Medical Center. Mouse Lewis lung cancer (LLC), human NSCLC (A549, H460, H1299, H1975, H358) and YFP-53BP1-HT1080 was maintained in DMEM with 10% fetal bovine serum at 37°C in a 5% CO_2_ humidified chamber. Human bronchial epithelial cell line BEAS-2B was maintained in RMPI 1640 medium (10% fetal bovine serum) at 37°C in a 5% CO_2_ humidified chamber as well. A549, H460, H1299, H1975, H358 and BEAS-2B cell was pre-treated with glucosamine at 1 hour before irradiation and further cultured for another 24 hours then switched to normal medium.

### SiRNA and transfections

Small interfering RNA (siRNA) oligonucleotide duplexes were designed against TG2 (sense, 5’-AAGGGCGAACCACCTGAACAA-3’ and antisense, 5’-TTGTTCAGGTGGTTCGCCCTT-3’) and TOPOIIα siRNA (purchased from Thermo Fisher, Catalog # AM16708). Plasmids and miRNA were transfected with Lipofectamine 3000 (Invitrogen) according to the manufacturer’s instructions. At different time after transfection, cells were subjected to further experiment.

### Irradiation

The ^60^Co γ-rays in Radiation Center (Faculty of Naval Medicine, Second Military Medical University, Shanghai, China) were applied for the irradiation exposure. After anesthetization with 10% chloralhydrate (350mg/kg), the mice were treated whole lung irradiation. All radiated animals received a single dose of 15Gy with a dose rate of 1Gy/min and were monitored up to 2 weeks post-irradiation. Cells were treated with 2, 4, 8Gy of γ-rays irradiation at a dose rate of 1Gy/min.

### Laser micro-IR

A 365-nm pulsed nitrogen laser (Spectra-Physics) was directly coupled to the epiflourescence path of the microscope (Axiovert 200M; Carl Zeiss) as described (43). Find the cell which placed on the cover slip two days before by using a Plan-Apochromat 63X/NA 1.40 oil immersion objective (Carl Zeiss, Inc). Then laser was used to generate DSBs in a defined area of the nucleus. At different time point, cells were fixed and subjected to immunofluorescence staining.

### Cell viability assay (CCK-8 assay) and clonogenic survival

A549 and BEAS-2B cells in good condition were seeded in 96-well plates in triplicate and after adherence treated with glucosamine. After 24h treatment, cell viability was measured by using a CCK-8 assay (Beyotime, Shanghai, China) according to the manufacture’s protocol.

Clonogenic survival was used to assess the potential of cell proliferation. Cells were calculated and seeded in the 6-wells plates, and then the pretreated cells were irradiated with 0, 2, 4, 8Gy. After incubated for 10 days, the plated were fixed with paraformaldehyde and stain with 1% methylene blue. The survival fractions were analyzed using a more target and one-hit model: f=1-(1-exp (-b*x)) ^c, where x is the dose in Gary and f is the survival fraction at dose x. b and c are the parameters of the survival curve.

### Apoptosis assay

After 24h post-irradiation, cell apoptosis were measured by double-staining with Annexin V-fluorescein isothiocyanate (Annexin V-FITC) and Propidium Iodide (PI) by Apoptosis Detection Kit (Invitrogen, Carlsbad, California, USA) and analyzed by flow cytometry (Beckman Cytoflex) according to the manufacturer’s instructions.

### Comet Assay

The DNA double-strand breaks of A549 cells were determined by ameliorating the aforementioned neutral comet assay. This part uses two-layer-agarose style, 1% normal melting agarose (NMA) as the base layer on the slide, 0.65% low melting point agarose (LMA) as the upper layer, suitable for smoothing and agarose adhesion, and The clarity of micrograms. Firstly, we prepared the slides by immersing the clean slides in a molten 1% NMA and wiping it immediately. All slides are pre-painted in advance to make sure they are thoroughly dry the next use (previous day). Next, the concentration of the single cell suspension prepared in ice-free Ca^2+^ and Mg^2+^-containing PBS was adjusted to 2×10^4^ cells / ml, and 0.4 ml of the solution was then immersed in 1.2 ml of LMA, 40°C water bath. Thirdly, 1.2ml of the cell suspension was mixed and rapidly pipetted onto the surface of the precoated slide. Fourth, once the solid solution, neutral solution (58.44 g NaCl, 5.584 g Na_2_EDTA and 0.61 g Tris) was dissolved in 500 ml of double distilled water, pH 8.2-8.5, Triton X-100 final concentration was 1% before use) Stay in the darkness. The slides were then gently soaked in TBE rinse buffer (0.744 g Na_2_EDTA, 10.902 g Tris and 5.564 g boric acid in 500 ml double distilled water PH8.2-8.5) and then incubated at 4 °C in the dark for 25 min at 25 V and 7 mA in fresh TBE. Fifth, the gel washed with ddH_2_O (double distilled water) was stained with PI (10μg/ml) for 20 minutes and then rinsed gently with ddH_2_O. Finally, all gels were examined by a fluorescence microscope (Olympus BX60) under a 10X objective. A total of 100 comet images in each slide were analyzed using special analysis software named CASP 1.2.3b2 (CASPlab, Wroclaw, Poland), which contained several features of DNA content, tail length, olive tail and tail.

### Western blotting and immunoprecipitation

Total proteins were obtained from cell lines using ProtecJETTM Mammalian Cell Lysis Reagent (Fermentas, Vilnius, Baltic, Lithuania) according to the manufacturer’s instruction. For nuclear and cytoplasmic protein, we using ProteinExt^®^ Mammalian Nuclear and Cytoplasmic Protein Extraction Kit. Briefly, cell pellets were resuspended(4×10^7^ cells/ml) in PBS by low-speed centrifugation (1000× g, 3min, 4°C), the pellets were lysed by 500 μl CPEB I and incubated in ice for 10mins, then added 55μl CPEB II and vortexed for 5s and incubated on ice for 1 min. After high-speed centrifugation (16000×g, 4°C) for 15min, the supernatant were collected as the cytoplasmic protein. Resuspended the pellets with 500 μl CPEB I, vortexed for 5s. Carefully discarded the supernatant, resuspended the pellets with 200 μl NPEB, incubated on ice for 30 mins and high-speed centrifugation(16000× g, 4°C) for 10mins. Collected the supernatant which was the nuclear protein. Then the samples were analyzed by western blotting with chemiluminescent detection as described elsewhere. For immunoprecipitation (IP), Cells were lysed in IP buffer (#9803,CST) and incubated overnight with pulled antibody-protein A beads(#9863,CST). The beads were washed with IP buffer and resuspended in 3X SDS Sample Buffer: (#7722,CST) for WB.

### Histopathology and immunohistochemistry

On 7 days after local lung irradiation, lung and tumor tissues were isolated, fixed and subjected to sectioning. Tissues were stained with H&E and antibodies for Ki67 (1:200; Cell Signaling Tech.), p65 (1:200; Cell Signaling Tech.), E-cadherin (1:200; Cell Signaling Tech.), Vimentin (1:200; Cell Signaling Tech.) and α-SMA (1:200; Cell Signaling Tech.). Five fields per section at ×200 magnifications were randomly selected per mouse, and two blinded pathologists independently examined 30 fields per group using Nikon DS-Fi1-U2 microscope (Nikon, Tokyo, Japan).

### Immunofluorescence analysis

We used an immunofluorescence assay to detect γH2AX foci (DNA double strand break marker), the subcellular location of 53BP1, TG2 and TOPOIIα. Briefly, A549 cells were seeded on 22X22mm^2^ cover glasses in 6-well plates at the concentration of 2X10^5^ per well. After different treatment, cells were fixed in 4% paraformaldehyde for 10min and permeabilized in 0.5% Triton X-100 for 10min. After blocked in serum, cells were stained with primary antibody (1:200) and then with the secondary antibody (1:1000). Cellular images were obtained using an Olympus BX60 fluorescent microscope (Olympus America Inc., Center Valley, PA, USA) equipped with a Retiga 2000R digital camera (Q Imaging Inc., Surrey, BC, Canada). Image Pro Plus (Media Cybernetics, Silver Springs, MD) were used to count the γH2AX foci per cell according to our previous studies, and at least 100 cells per group were counted.

### Patients samples and realtime PCR

Paired NSCLC and adjacent normal tissues resected surgically used for qRT-PCR were collected from 80 patients during operation at Shanghai Pulmonary Hospital (Shanghai, China). All the experiments were conducted with the informed consent of the patients and were approved by the ethics committee of Shanghai Pulmonary Hospital. Tissue array was performed by Zhuhao Tech., Shanghai. Total RNA was extracted from normal lung tissues and lung cancer tissues with TRIzol reagent, as described by the manufacturer (Invitrogen). The isolated RNA concentration was measured by Genequant pro (Biochrom Ltd, Cambiridge England). The RNA (1 μg) was applied to generate cDNA by means of PrimeScript™RT Master Mix (Takara, RR036A). The real-time quantitative PCR was performed by using SYBR Green Master Mix (Takara, RR420A) in StepOnePlus 96 Real-Time PCR System (Applied Biosystems). The average threshold cycle (Ct) of quadruplicate reactions was determined, and amplification was analysed by the ΔΔCt method. Gene expression was normalized to that of GAPDH. Real-time quantitative PCR with reverse transcription data were representative of at least three independent experiments. Primer sequences used to amplify human TG2 and TOPOIIα and GAPDH were as follows: TG2 forward: CCTGATCGTTGGGCTGAAG, TG2 reverse: TCGGCCAGTTTGTTCAGGTG; TOPOIIα forward: CCCACATCAAAGGCTTGCTG, TOPOIIα reverse: GATGTGCTGGTGCCCAAACC. GAPDH forward: AGCCACATCGCTCAGACAC, GAPDH reverse: GCCCAATACGACCAAATCC.

### Kaplan Meier Survival Analysis

We used an online tool Kaplan Meier (KM) plotter to analyze overall survival in lung cancer patients as described previously (44). Briefly, KM plot was obtained from the KM Plotter web-based (kmplot.com/analysis) curator, which includes relapse-free and overall survival data on 54,675 genes from 2437 lung cancer patients. In our analysis, patients were differed with TG2 expression, in combination with histology subtype, and radiotherapy. Populations were separated by median TG2 expression and plots were generated accordingly. From the KM database, three set of microarray Affymetrix probe 216183, 211573, 211003 were used in our analysis.

### Statistical analysis

Data were expressed at the means ± standard error of mean (SEM). Between group differences were tested using a one-way ANOVA. Two-group comparisons were performed using independent-samples Student’s t-test. P<0.05 was considered significant. All experiments were performed at least 3 independent times.

## Author contributions

Y.Yang. X.Lei. and Z.Liu.: study concept and design, carried out experiments, preparation of manuscript, obtain funding. K.Cao, Y.Chen: carried out experiments, data analysis, figures preparation. H.Qin, H.Qu, L.Liu. and Z.Liao: carried out experiments. J.Cai F.Gao Y.Yang: study design, obtained funding.

## Acknowledgement

This study was supported in part by the grants from National Natural Science Foundation of China (No. 31670861, No. 11635014, No. 11605289, No. 31700739)

---

**Figure S1.**
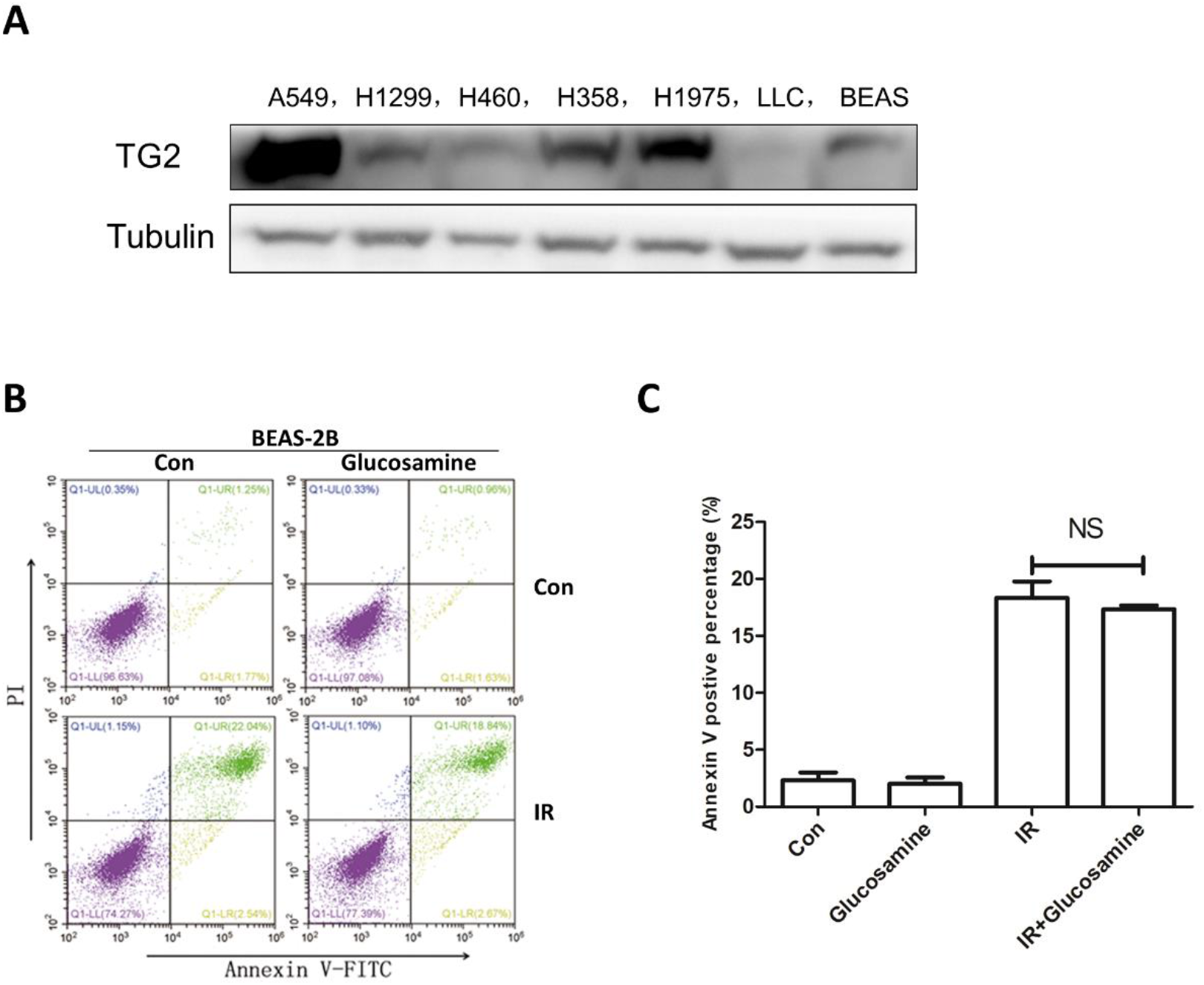
(S1A) Expression of TG2 in lung cancer cells including A549, H1975, H1299, H460, LLC and H358 cells, as well as normal BEAS-2B cells. (S1B, C) Representative images and column chart of flow cytometric analysis of BEAS-2B cell line against 8Gy irradiation with/without glucosamine (5mM) pretreated.

**Figure S2.**
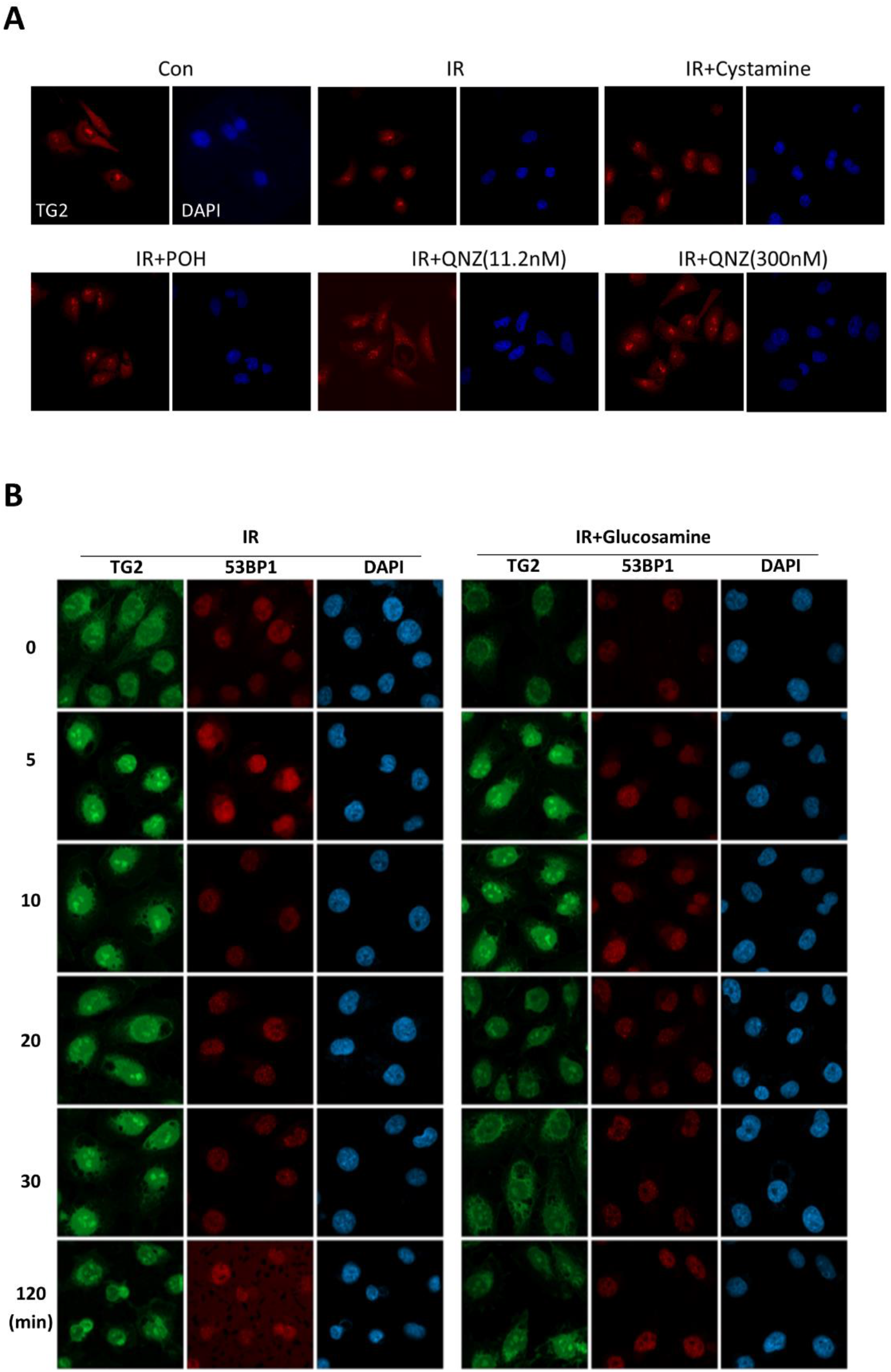
(S2A) Immunofluorescence of TG2 and DAPI in A549 cells exposed to IR with POH, cystamine or QNZ pretreatment. (S2B) Immunofluorescence of TG2 and 53BP1 in A549 cells exposed to IR with/without glucosamine (5mM) pretreated in delicate time point.

**Figure S3.**
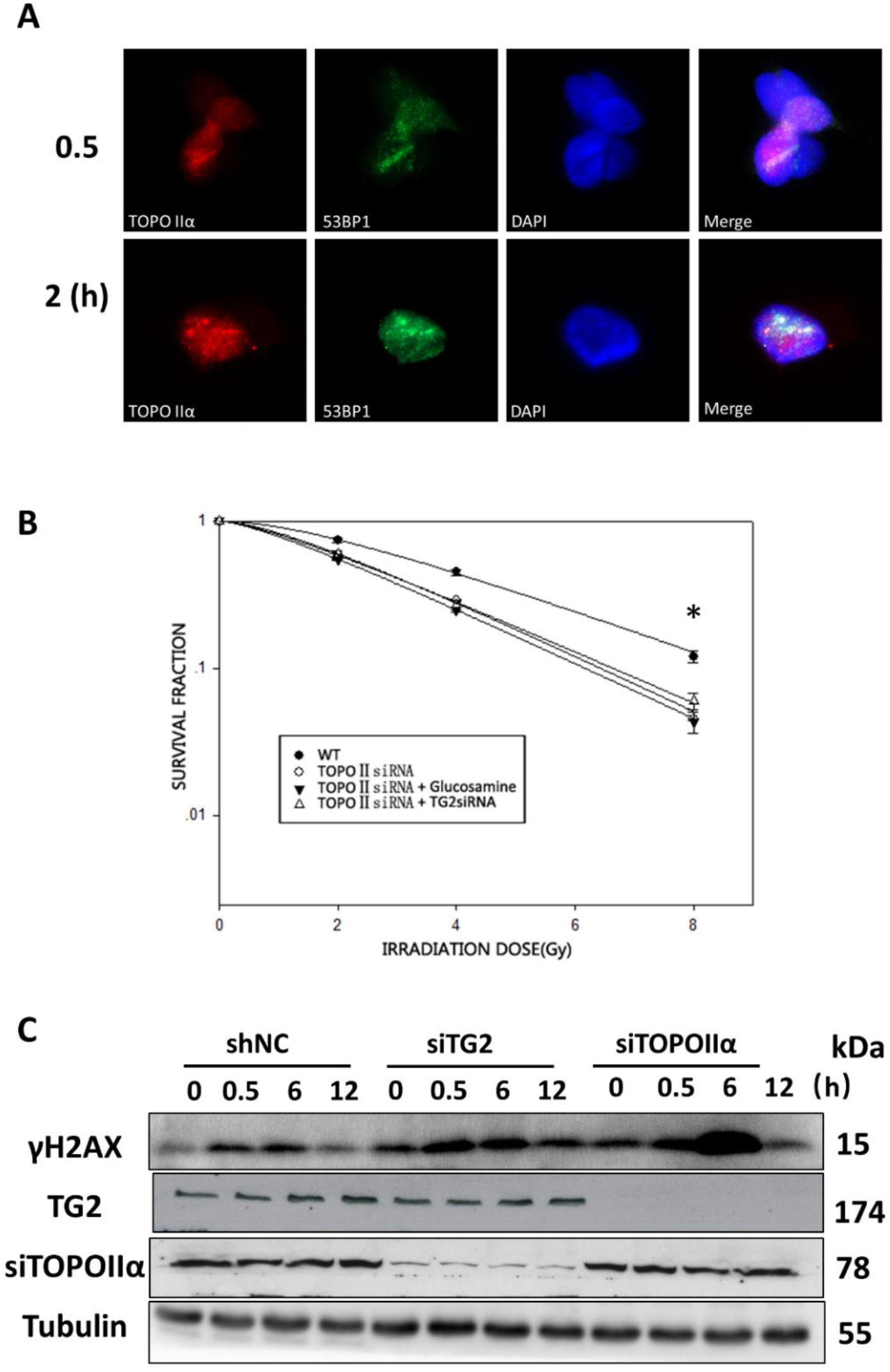
(S3A) Representative immunofluorescence of TOPOIIα and 53BP1 at DNA damage sites, in HT1080 cells following laser microirradiation. (n = 3 independent experiments). (S3B) A549 WT and TG2 KD cells transfected with TOPOIIα siRNA were analyzed for their colony forming ability against IR. (S3C) Expression of γ-H2AX in WT, TG2 knockdown or TOPOIIα knockdown A549 cell lines after IR.

**Figure S4.**
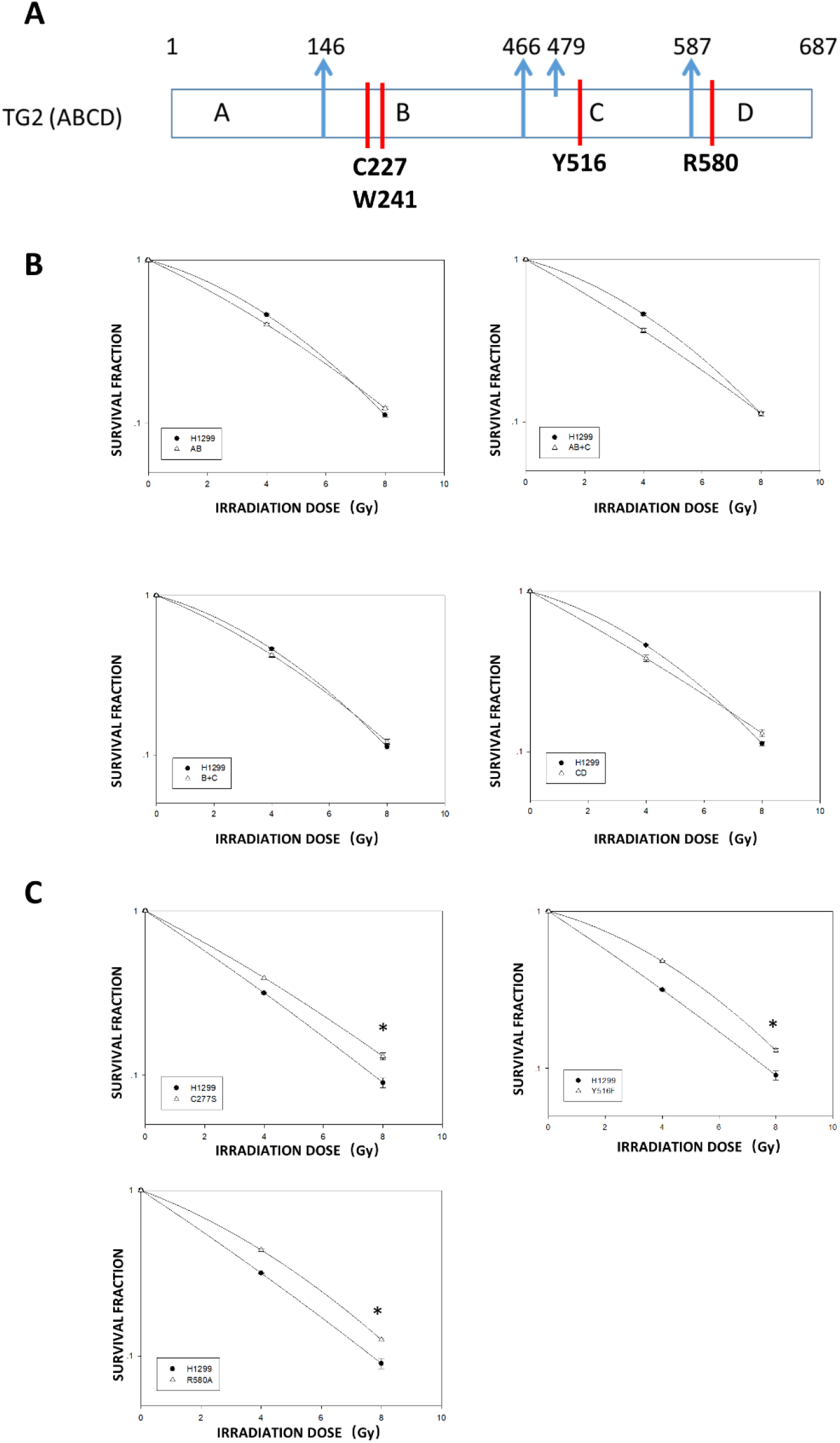
(S4A) Overlap of TG2 loss-function mutants and different fragment. (S4B) Survival of cells transfected with different TGM2 fragments plasmids (AB,AB+C, B+C,CD) were performed after irradiation. (S4C) Survival of cells transfected with different TG2 mutants (C277S, R580A, Y516F).

**Figure S5.**
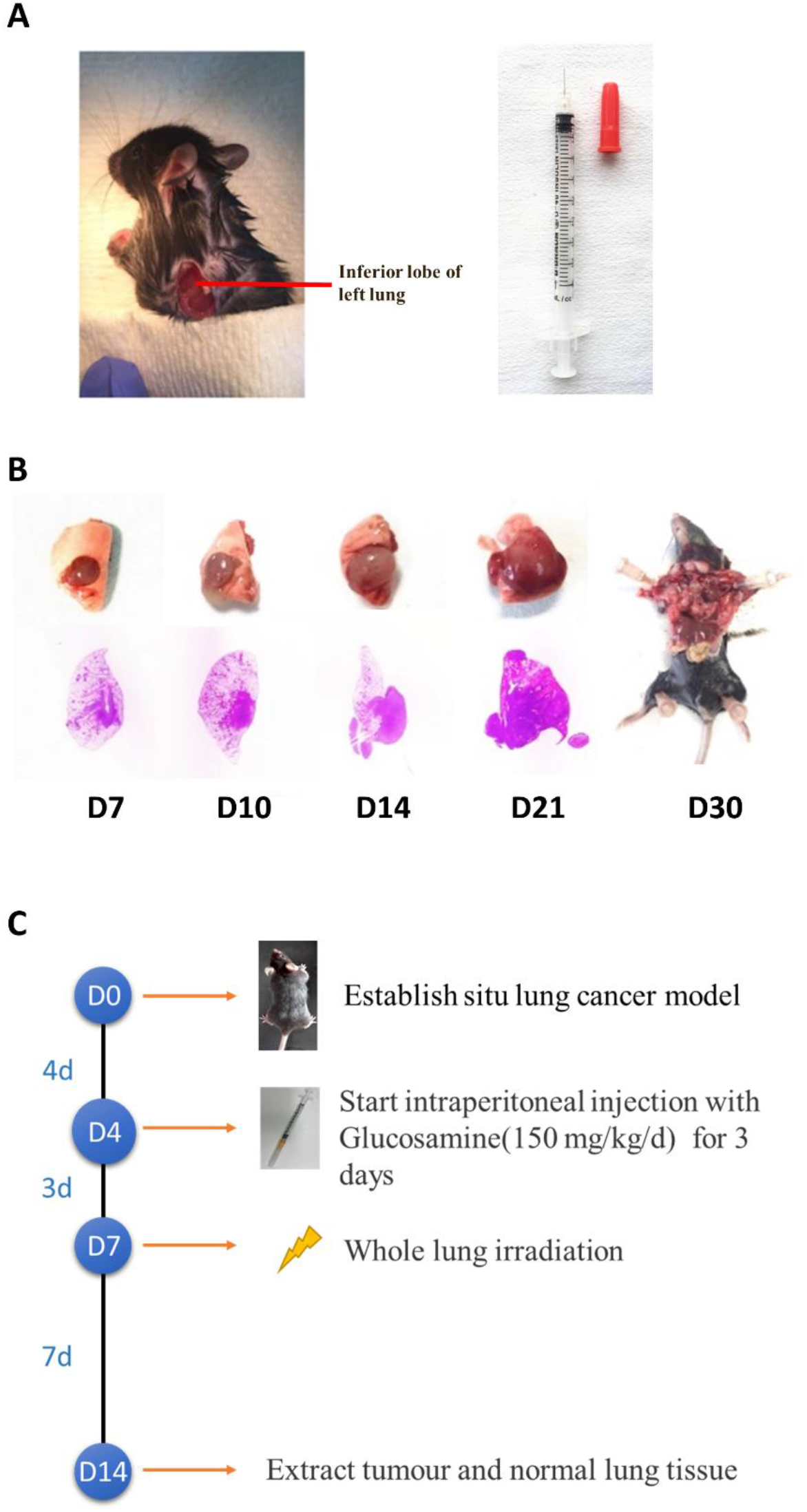
(S5A) The establishment of the in situ lung cancer model. (S5B) representative images of in situ lung cancer model and HE staining of each tumor. (S5C). time schedule of tumor implantation, radiotherapy and drug delivery.

**Figure S6.**
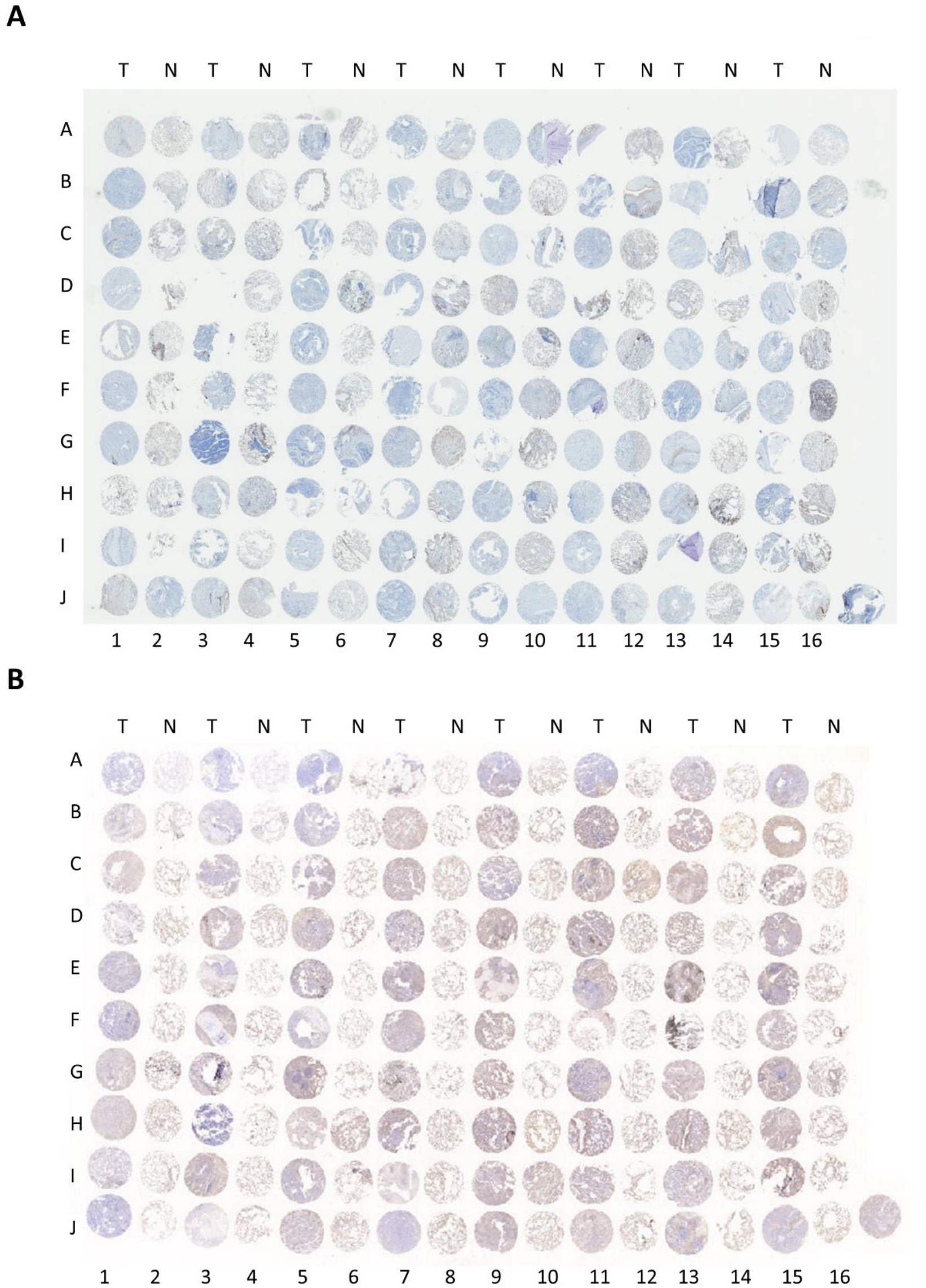
IHC staining of TG2 in two sets of lung cancer samples (total 160 pairs).

**Figure S7.**
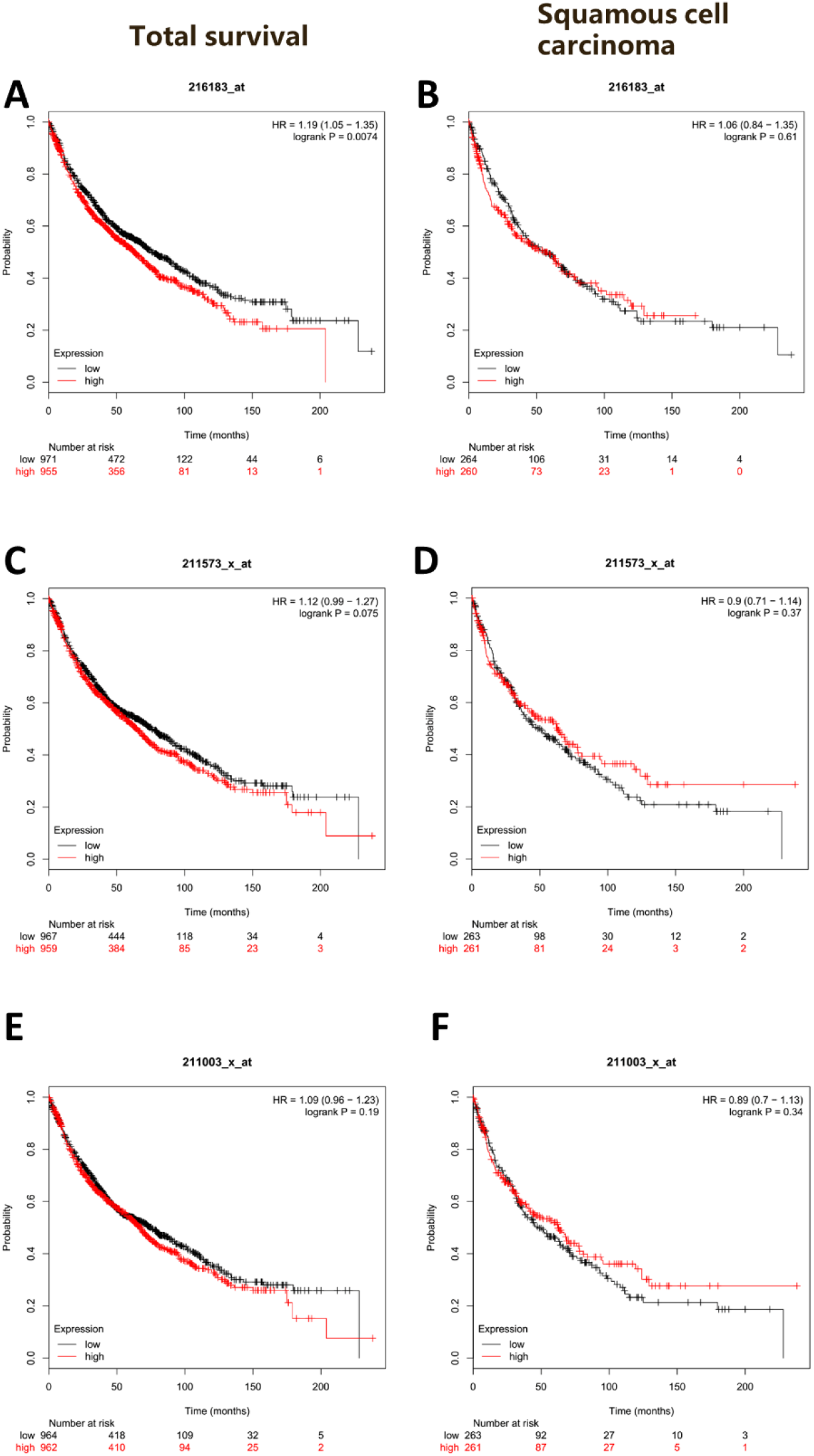
Kaplan-Meier survival curve showing little associations between TG2 IHC staining and overall survival of patients from lung squamous cell carcinoma and all patients from three sets of microarray.

**Figure S8.**
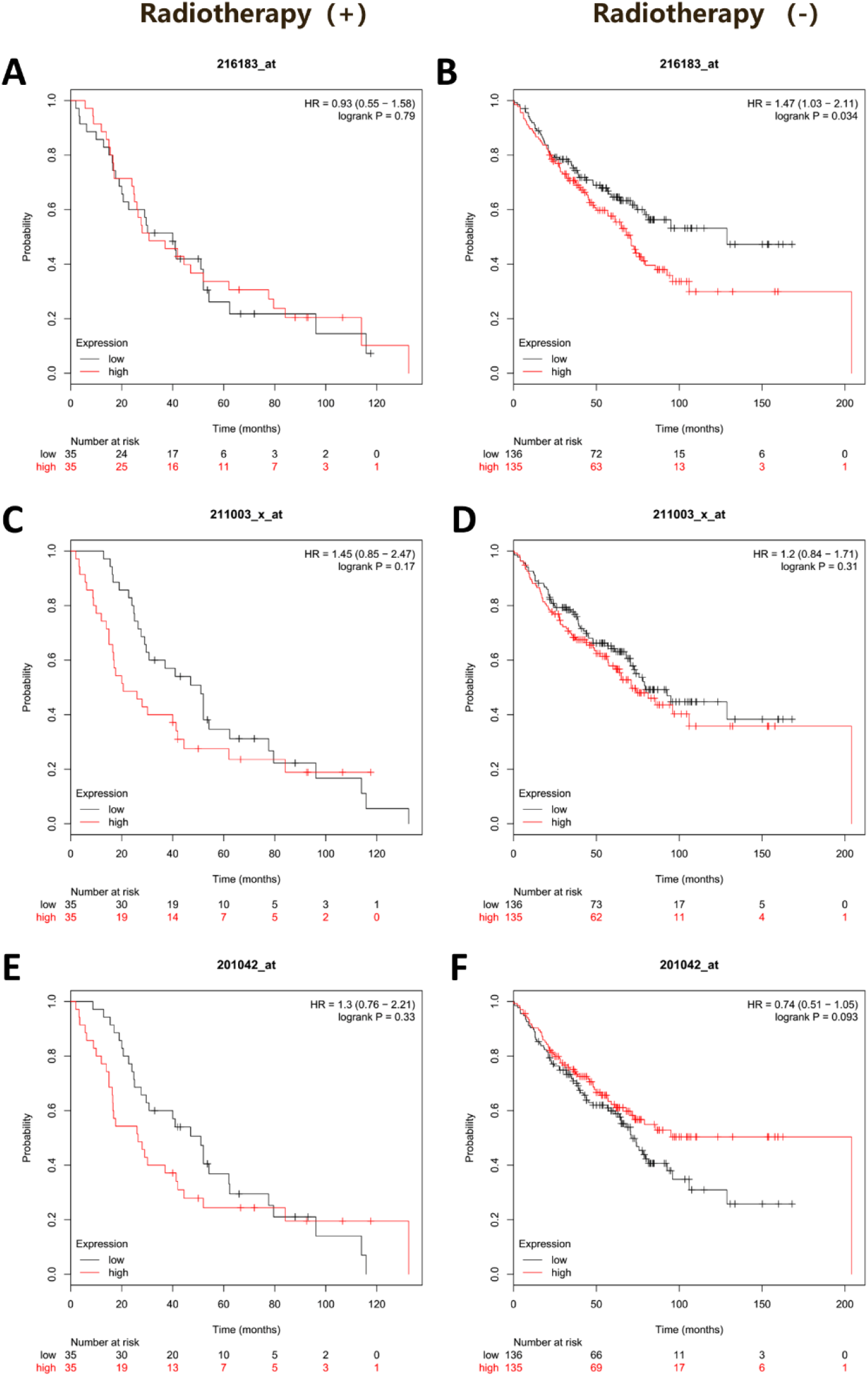
Kaplan-Meier survival curve showing no signifìncant association between TG2 IHC staining and overall survival of patients from lung cancer patients with and without radiotherapy.

**Table S2.** TG2 expression and clinical parameters

| Parameters | TG2 expression |  | Total<br>(n=80) | P value |
| --- | --- | --- | --- | --- |
|  | Low TG2 | High TG2 |  |  |
| Age (yr) |  |  |  |  |
| <60 | 20 | 21 | 41 | 1 |
| >60 | 20 | 19 | 39 |  |
| Sex |  |  |  |  |
| Male | 29 | 25 | 54 | 0.4739 |
| Female | 11 | 15 | 26 |  |
| Invasion |  |  |  |  |
| Yes | 8 | 9 | 17 | 1 |
| No | 32 | 31 | 63 |  |
| Histology |  |  |  |  |
| Adenocarcinoma | 21 | 28 | 47 | 0.1685 |
| No-Ade. | 19 | 12 | 31 |  |
| Adjuvant chemotherapy |  |  |  |  |
| Yes | 12 | 20 | 32 | 0.1101 |
| No | 28 | 20 | 48 |  |
| Smoking history |  |  |  |  |
| Yes | 18 | 11 | 29 | 0.1629 |
| No | 22 | 29 | 51 |  |

Note: see Table S1 in attached excel file.

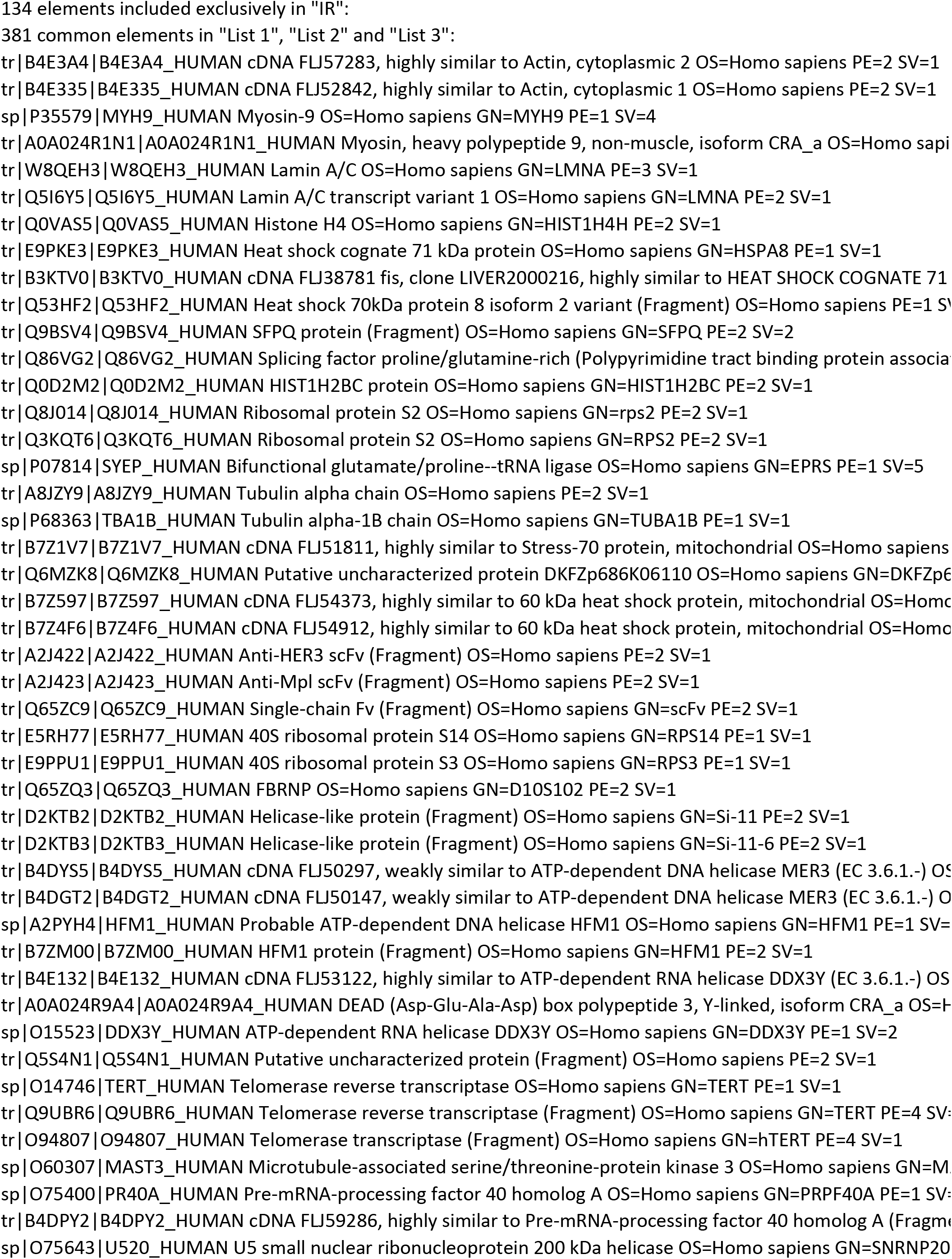
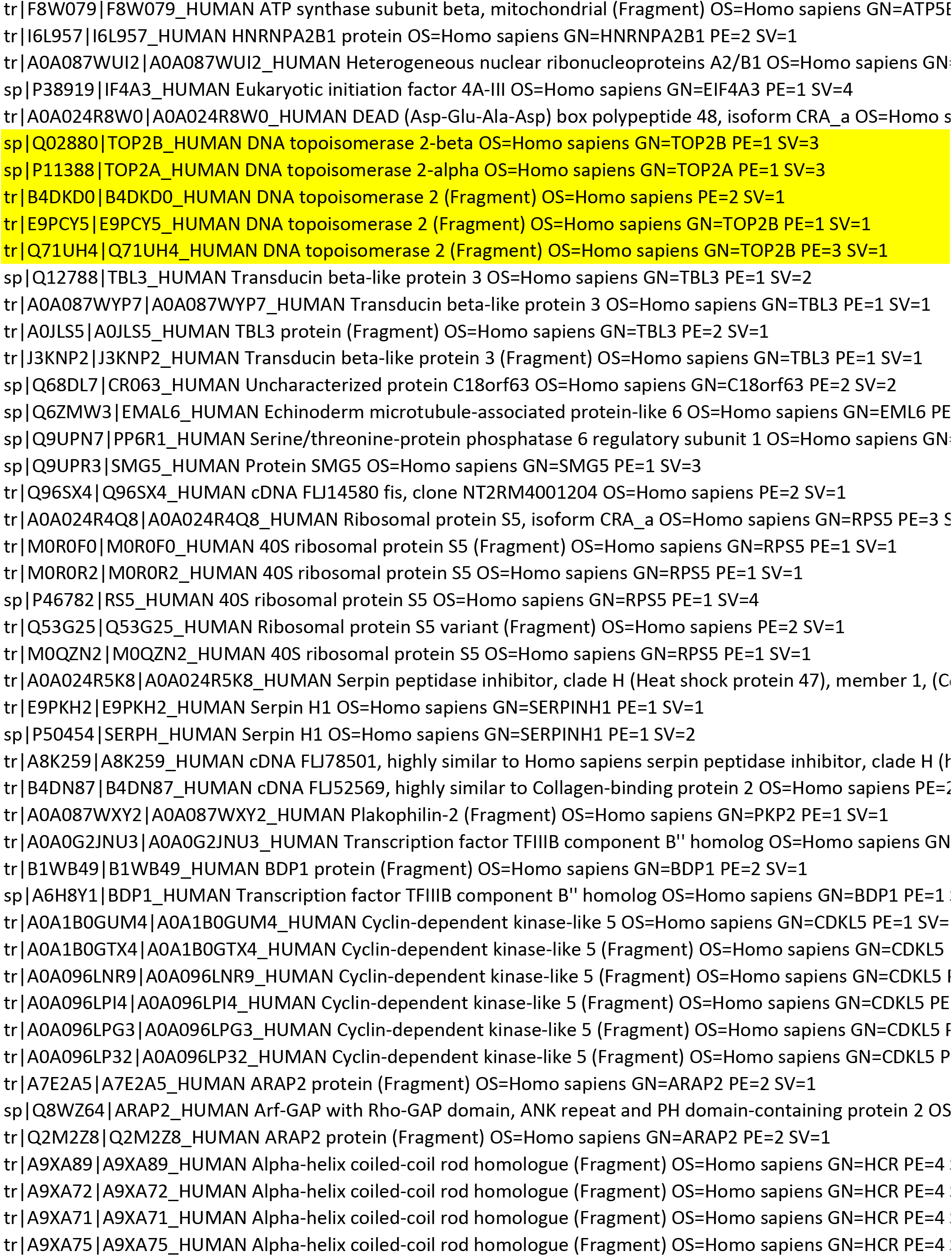
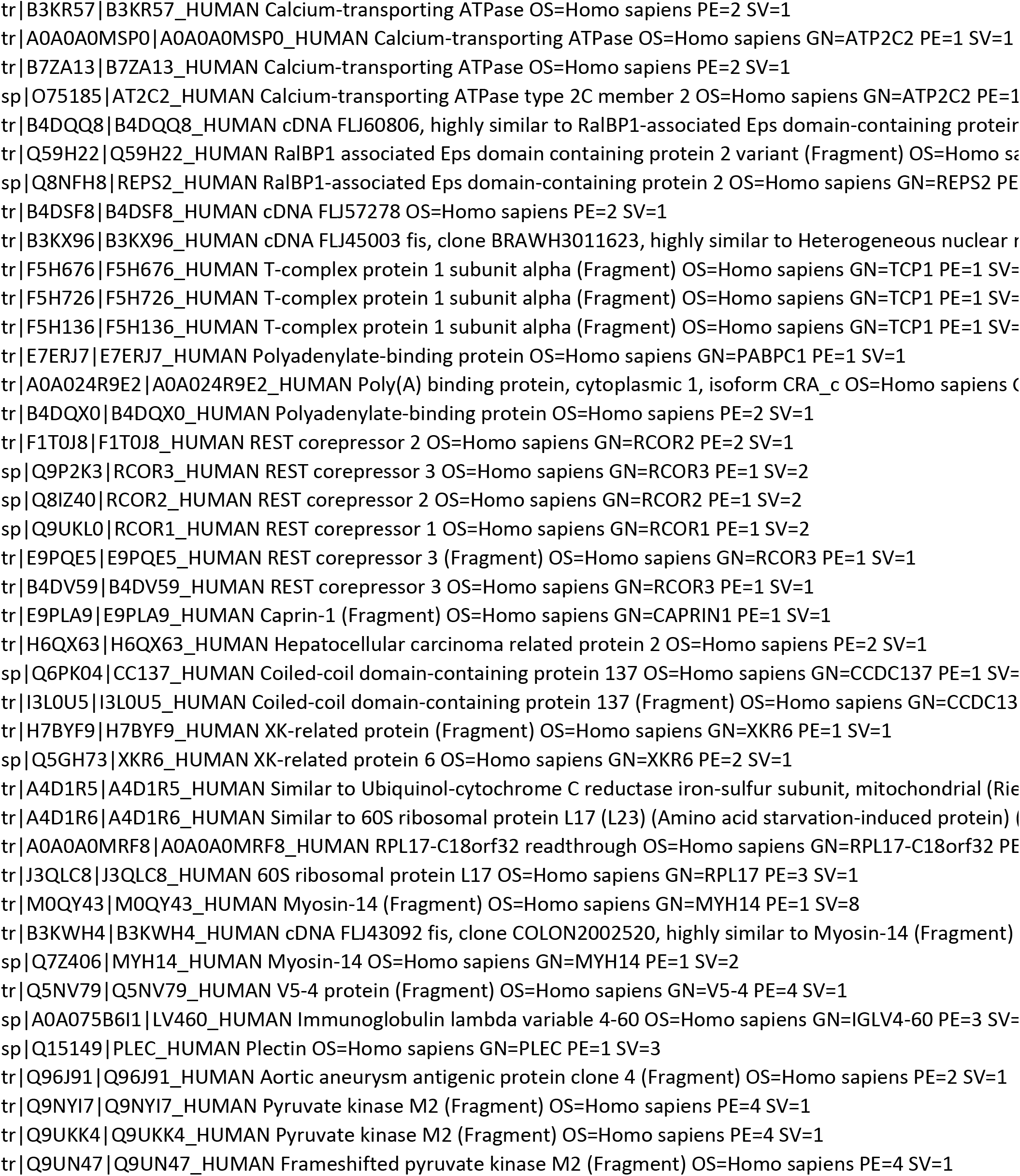
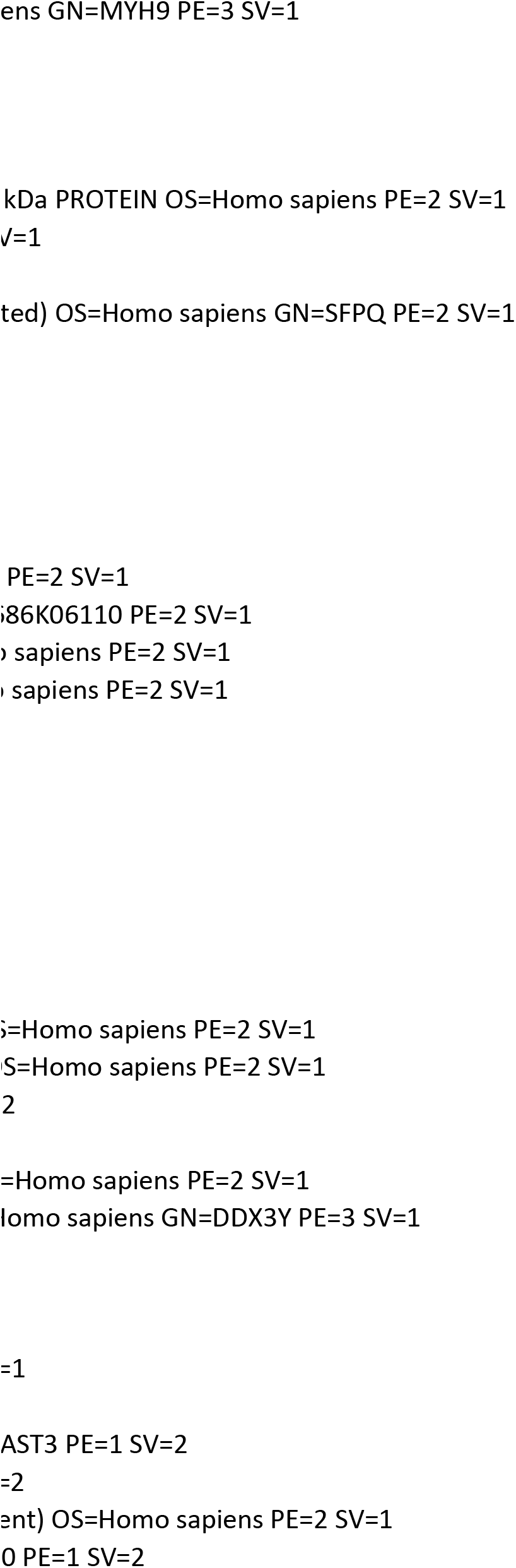
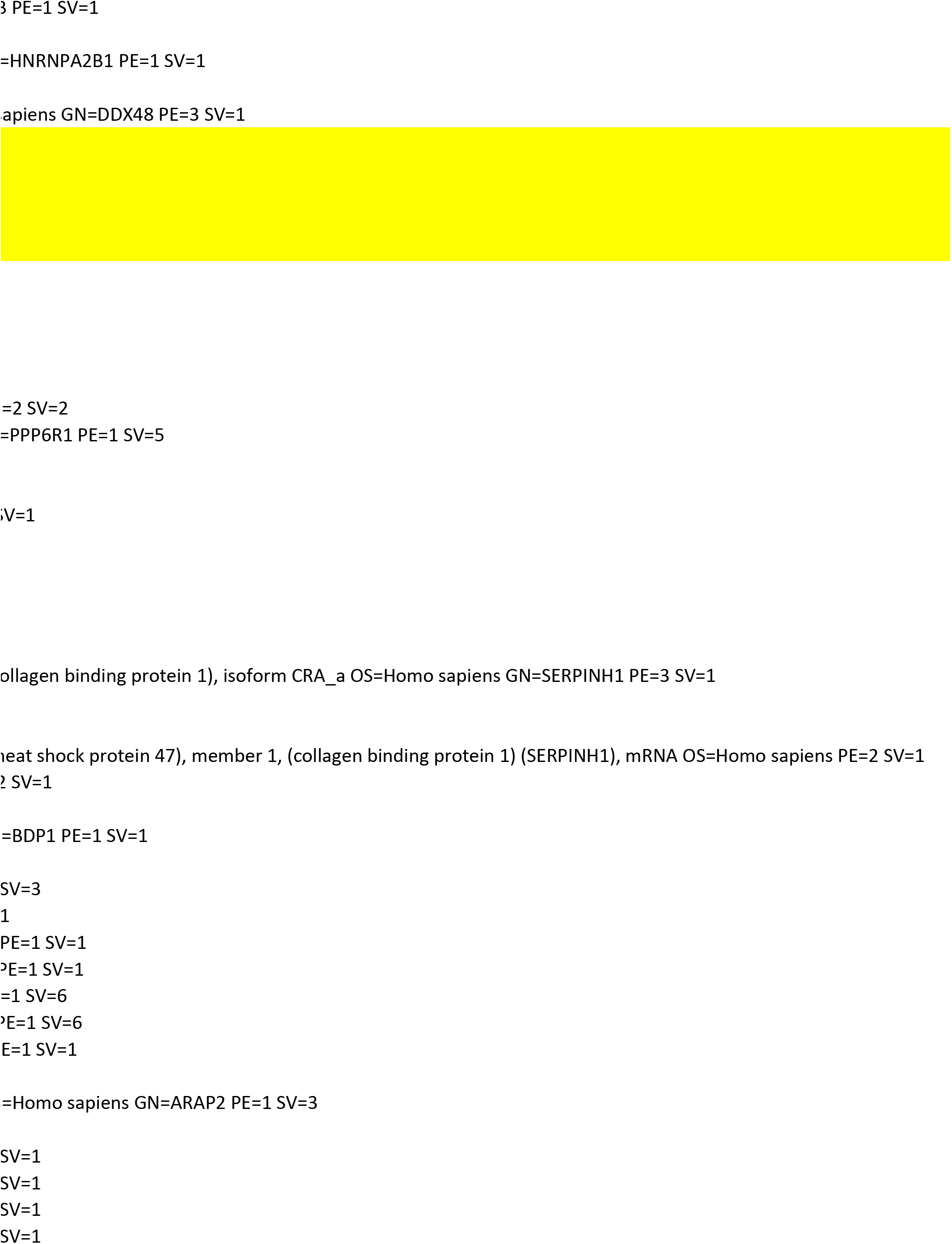
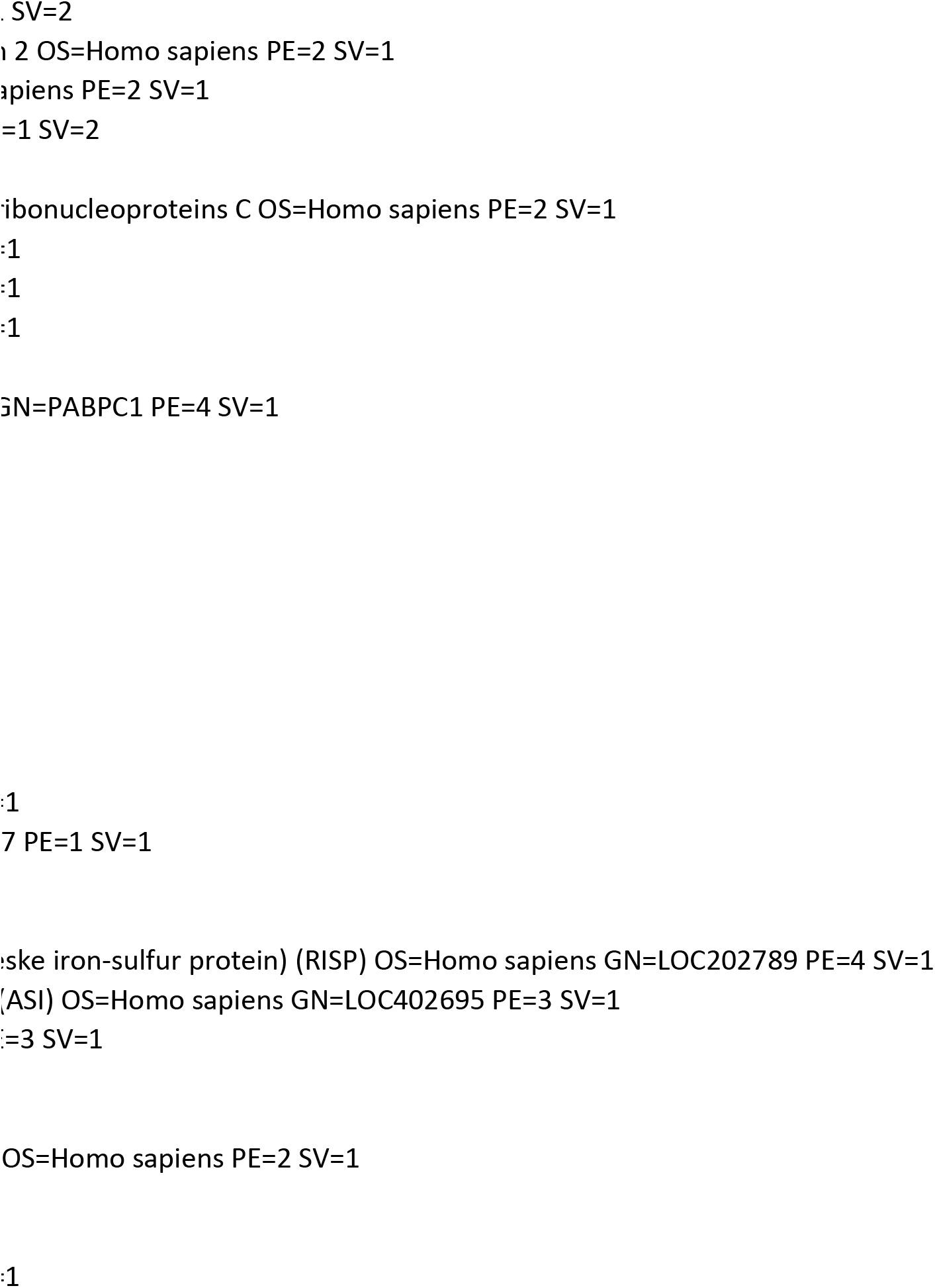

